# Matrix obstructions cause multiscale disruption in collective epithelial migration by suppressing physical function of leader cells

**DOI:** 10.1101/2022.04.28.489943

**Authors:** Ye Lim Lee, Jairaj Mathur, Christopher Walter, Hannah Zmuda, Amit Pathak

**Author notes:** To whom correspondence should be addressed:* Amit Pathak, Ph.D., One Brookings Dr., CB 1185, Saint Louis, MO 63130.

## Abstract

Cellular forces and intercellular cooperation generate collective cell migration. Pathological changes in cell-level genetic and physical properties cause jamming, unjamming, and scattering in epithelial migration. Separately, changes in microenvironment stiffness and confinement can produce varying modes of cell migration. However, it remains unclear whether and how mesoscale disruptions in matrix topology alter collective cell migration. To address this question, we microfabricated matrices with stumps of defined geometry, density, and orientation, which serve as obstructions in the path of collectively migrating healthy mammary epithelial cells. Here, we show that cells lose their speed and directionality when moving through dense obstructions, compared to those sparsely spaced. On flat surfaces, leader cells are significantly stiffer than follower cells, while dense obstructions lead to the overall softening of cells. In moving through dense obstructions, epithelial cells lose the sense of leaders and followers in their physical properties, migration phenotypes, and fluidity. Although Rac inhibition reduces obstruction sensitivity, loss of cell-cell cooperation and induction of leader-like phenotype via α-catenin depletion eliminates the effect of matrix obstructions on epithelial migration. Through a lattice-based model, we identify cellular protrusions, polarity, and leader-follower communication as key mechanisms for obstruction-sensitive collective cell migration. Together, microscale cytoskeletal response, mesoscale softening and disorder, and macroscale multicellular communication enable epithelial cell populations to sense topological obstructions encountered in challenging environments. These results reveal that cohesive, healthy populations are more obstruction sensitive than the dysfunctional, aggressive ones. The ‘obstruction-sensitivity’ could add to the emerging disease ‘mechanotypes’ such as cell stiffness and traction forces.

## Introduction

Collective systems are capable of complex motion due to motile entities within them that operate at varying lengths and time scales. A flock of birds can produce swirls and streams from altered behavior of small groups of birds or structural changes in bird wings; traffic patterns can dramatically change from minor disruptions around individual vehicles; collectively migrating cells can adopt varying modes depending on the type of cells or proteins that generate forces (1). Cell collectives, namely grouped epithelial cells, are important building blocks of all organs and tissue structures throughout the body. Their development and maintenance are crucial to the proper function of these organs (2). A key aspect of epithelial cells’ health and development is collective migration. During these processes, the migrating monolayer communicates signals from the leading edge throughout the trailing body to enable their navigation through heterogeneous environments (3). Migrating cell collectives have been physically described in terms of active matter that can transition across solid, fluid, or gas phases according to biochemical cues or the disease state of cells (4–6). Numerous studies have also shown that extracellular matrix (ECM) properties influence collective cell migration, from ECM stiffness and chemokine gradients to physical confinement and surface topography (7–9). Tissue and organ surfaces within the body (e.g., bone, lung, and skin) are not always smooth and have topological imperfections that can obstruct planar migration. It remains unclear how collectively migrating cells negotiate topological obstructions on matrix surfaces.

Collective cell migration is a multiscale process, regulated by mechanosensitive proteins, cell-ECM adhesions, cell-cell adhesions, and large-scale force propagation. In general, cells at the leading edge of migration, the leader cells, migrate outward such that biochemical and mechanical signals are efficiently propagated throughout the epithelial sheets via cell-cell adhesions (10, 11). This force transmission from the leader to the follower cells operates with such sensitivity that the entire monolayer can respond to mesoscale ECM stiffness gradients that change over just tens of microns (12). In this macroscale coordination, the dynamics of the cellular cytoskeleton regulate leader cell protrusions and the shape of all cells within the monolayer (13), while adherens junctions enable cell-cell communication and the collective stability and function of epithelia (14). Recent work has discovered that cell shape is an important indicator of the fluid-like behavior of the monolayer, defined by mesoscale phenomena known as jamming and unjamming (15). In general, unjamming refers to the transition to a fluid-like, pro-migratory state, characterized by polarized morphology and increased migration speed. By contrast, cell jamming typically entails a transition to a solid-like state, with rounded cell morphology and limited forward migration (15). The jamming-unjamming or solid-fluid transitions at cellular and population scales have been modeled as transitions between energy states, implemented in agent or lattice models (16). As the cell-cell contacts increase, the contractile energy of the cell minimizes, while the adhesive energy is maximized (17). Thus, the energy barrier between jammed and unjammed states can be viewed as a ratio between these two cellular energy levels (17). Taking into account random propulsion forces within the monolayer and cortical tension, cellular state transitions can be determined through the shape index of the cells, a ratio of the cell perimeter and area (18). This physical modeling technique allows for visualization of how microscale changes in parts of a cell or whole cells can yield varying outcomes on mesoscale and macroscale collective migration behaviors.

Just as changes within the cell, due to disease states such as asthma and cancer (19–21), affect the migration of the collective, so does the extracellular environment. Previous studies have shown that properties such as ligand density, geometry, and substrate stiffness all have pronounced effects on single-cell migration (22). In general, cells tend to generate higher cellular forces and, in turn, move faster on stiffer substrates or more confined environments (23, 24).

Ligand density has similar effects, such that increased density can lead to increased migration speed as well (25). Although the same ECM properties also influence collective cell migration, the mechanisms of force generation and signal propagation operate at multiple lengths and time scales, due to the size of the collective (26). As a result, microscale changes in the ECM can result in large-scale epithelial responses. Increasing the length of individual collagen fibers can result in sustained collective cell streaming due to increased cell-cell correlation (27). When physical defects (<50μm in size) are introduced in the basement membrane mimicking surfaces, healthy epithelial collectives can undergo disease-like epithelial-mesenchymal transition (EMT) and lateral invasion due to long distance propagation of degraded collagen-IV and cellular mechanoactivation (28). All of these effects occur due to effective communication among all cells within the monolayer. The function of a healthy epithelium or its dysfunction in cases of disease or injury necessitates the conversion of cellular level forces and signals into regional responses in the monolayer. Through development and disease, the ECM can degrade and remodel, which results in varied and heterogeneous regional topology within the matrix (29). As the ECM influences cell migratory behavior, it is important to further study how cell collectives sense and respond to topological heterogeneities, as well as how these signals are communicated throughout the monolayer on a cellular level. Since cell collectives have been shown to move similar to solids, fluids, or gases, depending on their cell type and environment (4, 6), it remains unpredictable how the migrating cell collectives would respond to topological obstructions in the matrix.

Altogether, collective cell migration phenotypes can change due to genetic mutations, protein conformations, cellular forces, multicellular cooperation, and environment properties – all of which span a wide range of length scales, from nanometers to millimeters. In this study, we broadly define microscale (<10μm) processes as those operating on the subcellular or single cell scale. Macroscale (>100μm) responses occur in the leader edge structure and net monolayer migration. Mesoscale properties occur in the intermediate length scale that covers small groups of cells or matrix structures in the 10-100μm range. To understand how mesoscale obstructions in matrix topology affect collective cell migration, we microfabricate Polydimethylsiloxane (PDMS) substrates with non-flat and stump-like geometries. Both the number and the orientation of the stumps were adjusted to study the effects of density and direction of obstructions on cell migration. We then utilized both bulk scale migration analysis, single-cell level stiffness via atomic force microscopy (AFM), live-cell tracking, and protein expression analyses to fully understand the sensing, transmission, and response of cells to the obstruction-filled environments. We also developed a computational model to understand how the mesoscale environmental changes result in microscale cellular modifications and macroscale obstructed collective migration. We discovered that dense matrix obstructions in the path of migrating healthy epithelia cause a slowdown in the forward migration, loss of directionality, reduced cellular fluidity, and cell-level softening, all of which result in disordered leading edges. This sensitivity to obstructions is differentially regulated by cytoskeletal remodeling and cell-cell cooperation. Overall, our results reveal that mesoscale matrix obstructions can cause multiscale physical changes in the collective migration of healthy cells. These observations may apply to other collective systems as long as individual entities within the system are sensitive to environmental cues and each other.

## Results

### Engineered matrix obstructions in the path of epithelial monolayers

To understand whether mesoscale obstructions in matrix topology can alter how epithelial cells migrate collectively, we use soft lithography to microfabricate Polydimethylsiloxane (PDMS) substrates with defined stumps. The dimensions of the stumps were kept constant at 100 μm by 25 μm (Fig. 1A), and their orientation was varied relative to the direction of cell migration. As a matter of terminology, in this study, we will refer to these topological features as ‘obstructions’ when discussed from the standpoint of cells, and as ‘stumps’ in the context of matrix properties. We varied spacing between individual stumps such that ‘sparse’ and ‘dense’ obstructions are 50 and 150 μm apart, respectively. We also varied the orientation of stumps relative to the direction of cell migration, such that they present either short obstructions of 25 μm in the case of parallel orientation, or long obstructions of 100 μm in the case of perpendicular orientation (Fig. 1B).

**Figure 1:**
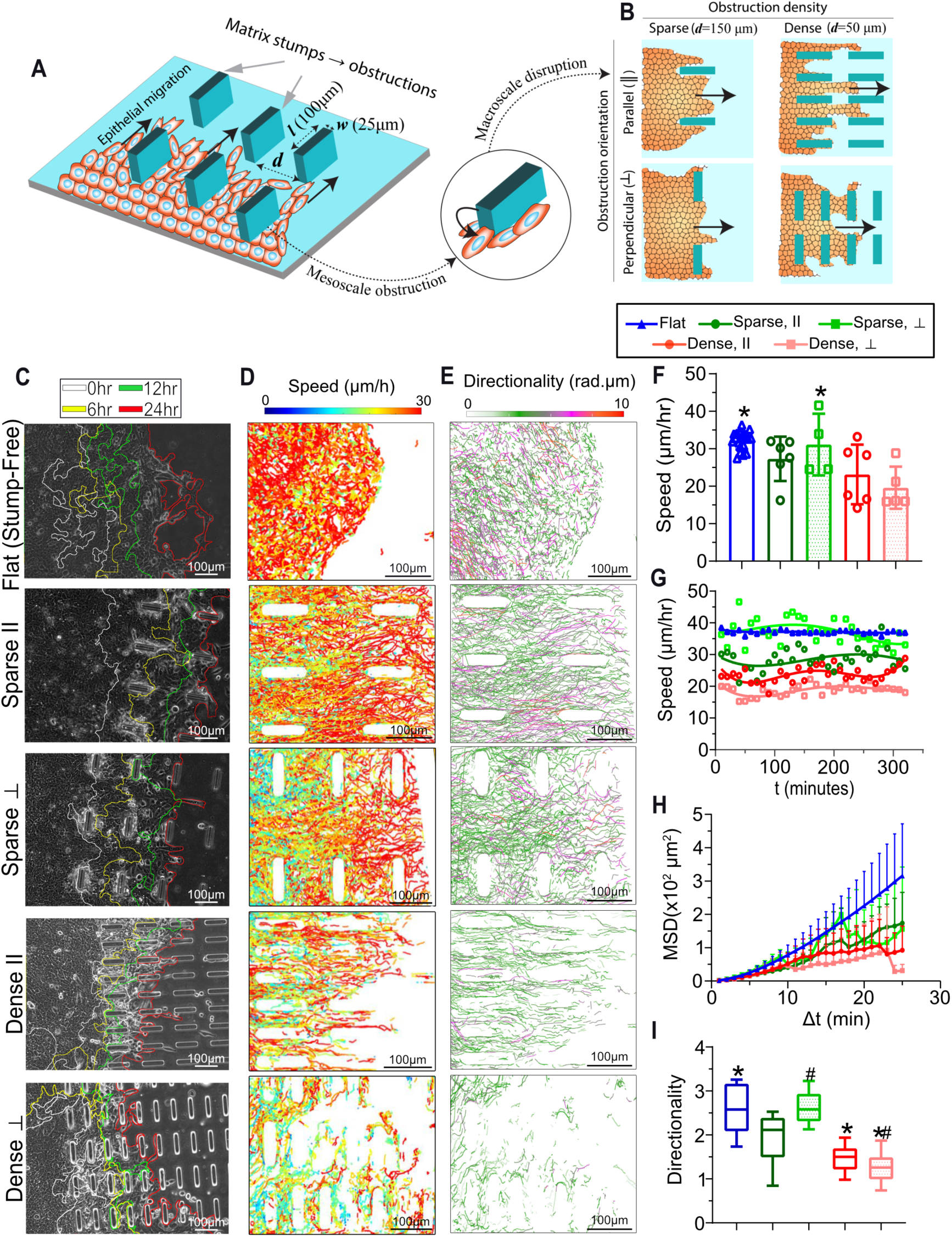
Dense obstructions reduce collective cell migration speed and directionality. **(A)** Schematic describing collective epithelial migration on engineered substrate with rectangular stumps of dimensions *l* = 100μm and *w* = 25μm, which serve as mesoscale matrix obstructions. **(B)** Varying spacing and orientation of stumps generate four different conditions for obstructions in the path of collective cell migration. **(C)** Representative phase contrast images of migrating MCF10A epithelial cells in different substrates at final timepoint (24hr after cell seeding), with leading edges annotated at previous timepoints in 6hr intervals. Migration trajectories of individual cells within epithelial monolayers are plotted with color coding for time-averaged **(D)** speed and **(E)** directionality. **(F)** Migration speed averaged over all cells per condition over 3 days of migration. Error bars = SD. **(G)** Temporal variation in average migration speed for each substrate condition; points represent average across all cells for a specific timepoint, lines are polynomial fits across timepoints. **(H)** Plot of mean squared displacement (MSD) versus duration of time interval. Error bars = SEM. **(I)** Directionality of cell migration averaged over all cells per condition over 3 days of migration. Error bars = SD. Significant difference annotated as *p<0.05 for comparison against the sparse condition, and #p<0.05 for comparison against the parallel condition. N ≥ 6 fields of view in all plots.

These dimensions were chosen such that the whole cell body could interact with stumps. The substrate was designed such that epithelial cell monolayer is first seeded in an open space, without stumps, and allowed to migrate outwards before running into obstructions presented by the stumps (Fig. S1 and supplementary Fig. S1; detailed in Methods). After coating the PDMS substrates with fibronectin, we cultured MCF10A human mammary epithelial cells. Through live-imaging, we studied cell interaction with stumps and various collective migration phenotypes over time through these varying obstruction courses. For initial understanding, these measurements are first compared against a control condition of PDMS substrates with stump-free ‘flat’ space; however, the key focus of the study is to compare the effects of sparse versus dense and parallel versus perpendicular obstructions on collective cell migration.

### Multiscale disorder in obstructed collective cell migration

To measure the response of collectively migrating cells to obstructions presented by matrix topology, we performed time-lapse imaging over 24hr and analyzed trajectories of individual MCF10a cells within the epithelial monolayer. In these cell migration videos, we observed that epithelial monolayers moving through denser obstructions, particularly those in a parallel orientation, had more ruffled leading edges and overall shorter distance traveled, compared with sparse-stumps or stump-free (flat) substrates. As shown in Fig. 1C, the epithelial monolayer migrating through open space or sparse obstructions progressed over time without significant backward retraction. However, when migrating through dense obstructions, the leading edges (tracked at 6hr intervals) intersected with each other, indicating their dynamic frontward and backward movement as they navigated around each obstruction. As a result, the leading edge became chaotic and ruffled when moving through many obstructions, particularly in the dense condition (Fig. 1C, each time frame visualizations provided in supplementary Fig. S2).

Macroscale disruption in the leading edge integrity caused by dense obstructions likely alters cell-scale migration behaviors. To investigate this, we tracked and plotted the speed of all cells that form the collective monolayer. As visualized in Fig. 1D, on stump-free, open substrates, cells were homogenously fast, regardless of their distance from the leading edge into the colony. However, the presence of obstructions (sparse and dense) led to more heterogeneous and overall slower migration speeds of cells within the monolayer, with faster cells at the leading front. Plotting the average migration speed across samples (Fig. 1F) confirmed these observations of more heterogeneous (see bigger error bars in Fig. 1F), yet slower average migration speeds in the case of dense obstructions, compared to sparse or no obstructions.

Throughout the full 30hr duration of tracking, cell migration speed in all conditions remained steady (Fig. 1G). Among these tested substrates, cell migration was the fastest on substrates with sparse-perpendicular stumps or those without any stumps. In other words, cell migration was the fastest when the contact area of obstructions in the direction of migration was the smallest. When sparse stumps were oriented parallel to the direction of migration, the migration speed slowed. Dense stumps led to the slowest migration overall, with similar speeds for both orientations (Fig. 1F).

In previous work, it has been shown that epithelial cell migration is most efficient when they adopt a fluid-like phenotype (21). Since obstructions reduce individual cell speeds in our data, we hypothesized that the physical impact of obstructions on cells likely manifests itself in an overall reduction in the fluidity of the system. To understand this, we measured mean square displacement (MSD) relative to varying time intervals averaged for all cells (Fig. 1H), as done previously (21). Here, higher slopes indicate more fluidic cell migration on stump-free flat surfaces and lower fluidity through sparse and dense obstructions. To better understand how the loss of collective cell fluidity arises, we measured the directionality of individual cells, defined as a product of direction autocorrelation and cell displacement (see Materials and Methods).

According to this definition, higher directionality indicates more persistent and ordered migration towards the leading edge. We found that cell migration directionality reduced with increasing density of obstructions (Fig. 1F). In the case of sparse obstructions, there was a gradient of higher directionality towards the leading edge cells. However, in dense obstructions, we noted that the spatial distribution of directionality was somewhat homogeneous across the monolayer. Together, these results show that mesoscale matrix obstructions reduce microscale directionality and migration speed of cells within the epithelial monolayer, which results in macroscale loss of fluidity and chaotic leading edge shapes.

### Cellular softening around dense obstructions

According to our results, matrix obstructions slow collective cell migration through mesoscale disorder. Do microscale mechanical properties of cells change as they navigate these topological obstructions? Although cell-level tractions in epithelial populations have been connected to varying jamming and unjamming phenotypes of collective cell migration (5, 20, 21), it remains unknown whether cell stiffness depends on matrix features. In the context of matrix obstructions, it is a particularly relevant question because of the direct cell-body interaction with the obstructions and the already known softening of cells moving through confining environments (30). To our knowledge, cell stiffness in collectively migrating monolayers has not been measured. We assessed changes in cell stiffness through atomic force microscopy (AFM).

Briefly, substrates were loaded onto the AFM stage so that the stumps ran parallel to the AFM cantilever to ensure that the AFM probe interacted with the cells without colliding with the stumps (Fig. 2A). We used AFM probes with spherical tips of 4.5μm diameter and indented cells over 0.25μm depth to acquire force-displacement curves, which were processed using a modified Hertz model to calculate microscale cell stiffness (details in Methods). Testing began at the leading edge of the monolayer, and full runs were taken in 100μm increments until the region of the monolayer before the first stump was reached. Each section was selected such that measurements were taken as close to the stumps without the cantilever touching the stumps and thereby skewing the stiffness measurements.

**Figure 2:**
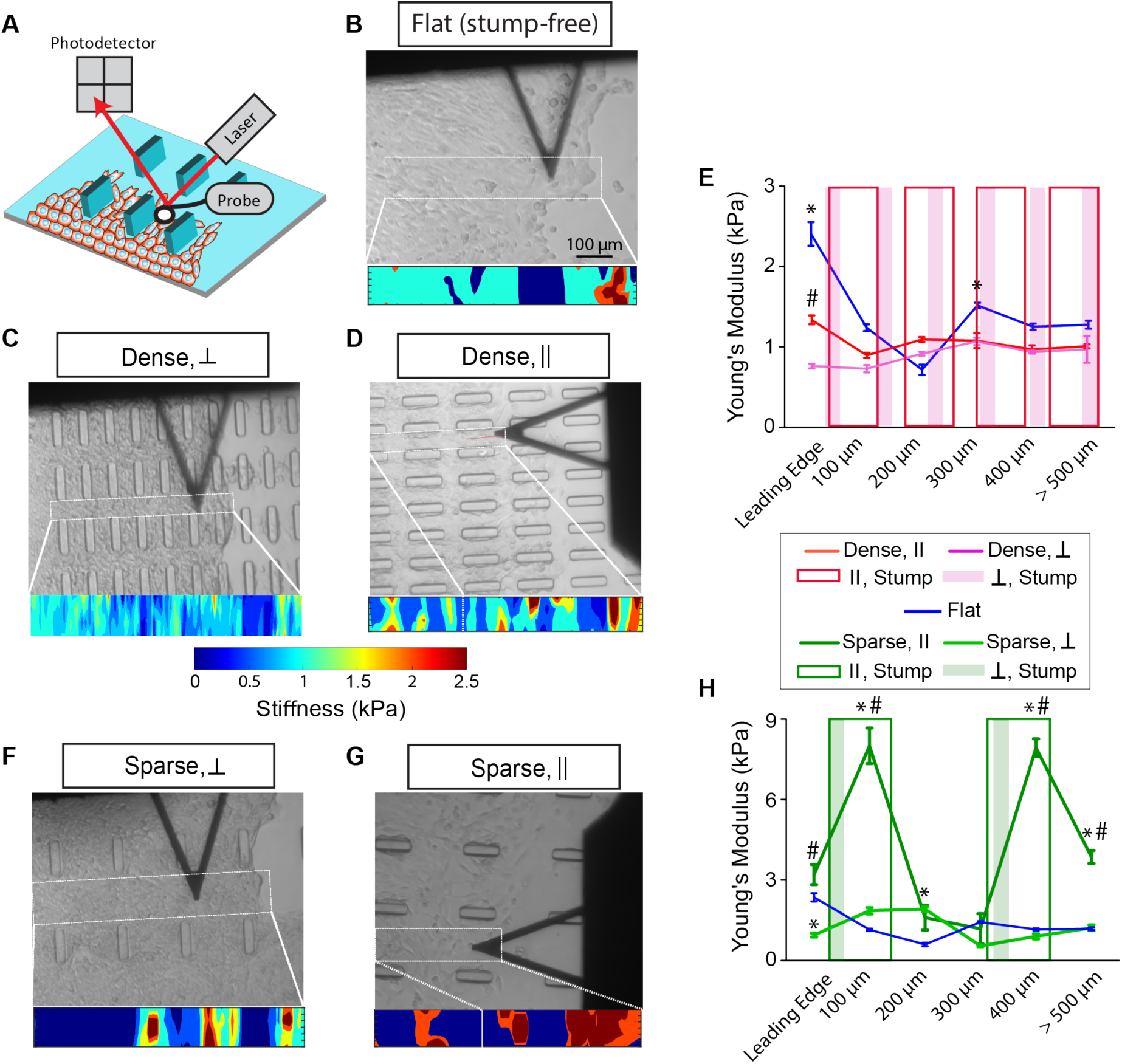
Cell stiffness responds to mesoscale obstructions. **(A)** Representative diagram of how cell stiffness was determined using AFM (see Materials and Methods). **(B-D)** Representative heat maps of stiffness scans along with corresponding brightfield images for **(B)** flat, **(C)** dense-perpendicular, and **(D)** dense-parallel conditions. White boxes and bars indicate the corresponding region where the heatmap aligns with the cell image. Scale bar = 100 *µ*m. **(E)** Average cell stiffness by distance from the leading edge for flat, and dense obstructions with both parallel and perpendicular orientations. Shaded bars indicate the location of stumps in their respective conditions. (F,G) Representative heat maps of full stiffness measurement runs with corresponding brightfield images for **(F)** sparse-perpendicular and **(G)** sparse-parallel stumps. White boxes and bars indicate the corresponding region where the heatmap aligns with the cell image. Scale bar = 100 *µ*m. **(H)** Average cell stiffness by distance from the leading edge for flat and sparse obstructions with both parallel and perpendicular orientations. Significant difference annotated as *p<0.05 for comparisons against flat substrates, and #p<0.05 for comparison between parallel and perpendicular conditions. Error bars = SEM. N ≥ 3 independent samples.

As a control, we first measured the stiffness of cells that form the epithelial monolayer on flat substrates by scanning from the leading edge into the monolayer. As shown in the representative monolayer and the spatial AFM scan, cells at the leading edge were stiffer compared to the cells within the monolayer, with average Young’s moduli reducing from approximately 2.8 to 0.5 kPa (Fig. 2B), i.e., leader cells were much stiffer than follower cells embedded within the epithelial monolayer. Although this variation in leader-follower cell stiffness has not been shown before, it is not entirely surprising given the known rise in mechanosensitive proteins, phosphorylated myosin light chain (pMLC) (9) and yes-associated protein (YAP) (31), towards the leading edge. To understand the effect of matrix obstructions on cell stiffness, we performed AFM on cell monolayers that had already migrated into dense stumps. According to the representative spatial maps for cells around dense stumps, cell stiffness was generally lower compared to the flat substrates (Figs. 2C,D). Going from the leading edge into the monolayer, cell stiffness was relatively homogenous and consistently lower in the case of dense-perpendicular obstructions (Fig. 2C), compared to more distinct patches of soft and stiff cells in parallel obstructions (Fig. 2D). For these three substrate configurations, we repeated AFM measurements for at least 3 different samples and averaged the cell stiffness, which was binned relative to distance from the leading edge, as plotted in Fig. 2E (location of the stumps is denoted in the background). On average, cell stiffness at the leading edge was the highest in the epithelial monolayers on control (more intact) flat substrates (~2.5 kPa) and reduced by almost half around dense parallel obstructions (~1.4 kPa), with even further reduction in case of dense perpendicular obstructions (~0.8 kPa). These results (Figs. 2B-E) show an overall cell softening, particularly at the leading edge, when epithelial cells are forced to navigate dense matrix obstructions.

Next, we sought to test changes in the stiffness of cells moving collectively through sparse stumps, an intermediate presentation of obstructions. In sparse-perpendicular obstructions, as shown in the representative AFM map in Fig. 2F, we observed patches of high cell stiffness (~2.5 kPa), but overall cell stiffness across the epithelial cell monolayer seemed lower than that on flat substrates. When we repeated these measurements in sparse-parallel obstructions, heterogeneity in cell stiffness across the monolayer was further accentuated, with ‘stiff patches’ of much higher cell stiffness compared to any other condition (Fig. 2G). Given that dense obstructions softened cells, higher cell stiffness in the case of intermediate stump density, although only in the case of parallel orientation, was a surprising result. To confirm these trends and find a potential explanation for these results, we repeated these measurements 3 more times on different monolayers. The overly stiff patches were localized in epithelial regions between parallel stumps. Upon plotting cell stiffness as a function of distance from the leading edge and relative to the location of stumps, the average cell stiffness sharply increased, to approximately three-fold, in regions between the parallel stumps. In all other regions, cell stiffness values were similar to those in flat or sparse-perpendicular substrates (Fig. 2H).

According to these results, although dense obstructions broadly softened the cells within the epithelial monolayer (Figs. 2C-E), sparse-parallel stumps led to a complex effect on cell stiffness (Fig. 2G, H). Overall, there are two key takeaways from these AFM measurements: (1) on flat substrates, leader cells are significantly stiffer than follower cells; (2) dense obstructions soften the leader cells and thus make them behave like follower cells.

### Loss of cell shape fluidity due to matrix obstructions

As epithelial colonies move through and around matrix obstructions, individual cells within the colony lose directionality and speed in their migration (Fig. 1), along with general softening and mesoscale spatial heterogeneity in cell stiffness relative to distance from the leading edge and density of obstructions (Fig. 2). Following these trends down the length scale – from the leading edge roughness at the scale of hundreds of microns to migration and stiffness at the scale of tens of microns – we next sought to measure cytoskeletal and cell-level changes due to obstructions. Given the known importance of actin polymerization in protrusions that propel migration (32, 33) and the actin-rich cytoskeleton that provides structural strength to the cell (34), we stained with phalloidin to measure F-actin intensity. We found higher actin expression near the leading edge on all substrates, albeit with some obstruction-dependent distinctions (Fig. 3A). In denser obstructions, we noted that areas of high actin intensity were smaller and localized closer to the leading edge. By contrast, cell monolayers migrating through sparse stumps showed broader regions of high F-actin intensity near the leading edge. After averaging across multiple samples, we found a significant reduction in actin expression in dense obstructions compared to sparse obstructions (Fig. 3B). We noted the highest actin expression was in sparse-parallel obstructions. Thus, the unexpectedly high cell stiffness in this condition (discussed above in Fig. 2) could potentially be due to the increased actin expression (Fig. 3B), which remains unexplained.

**Figure 3:**
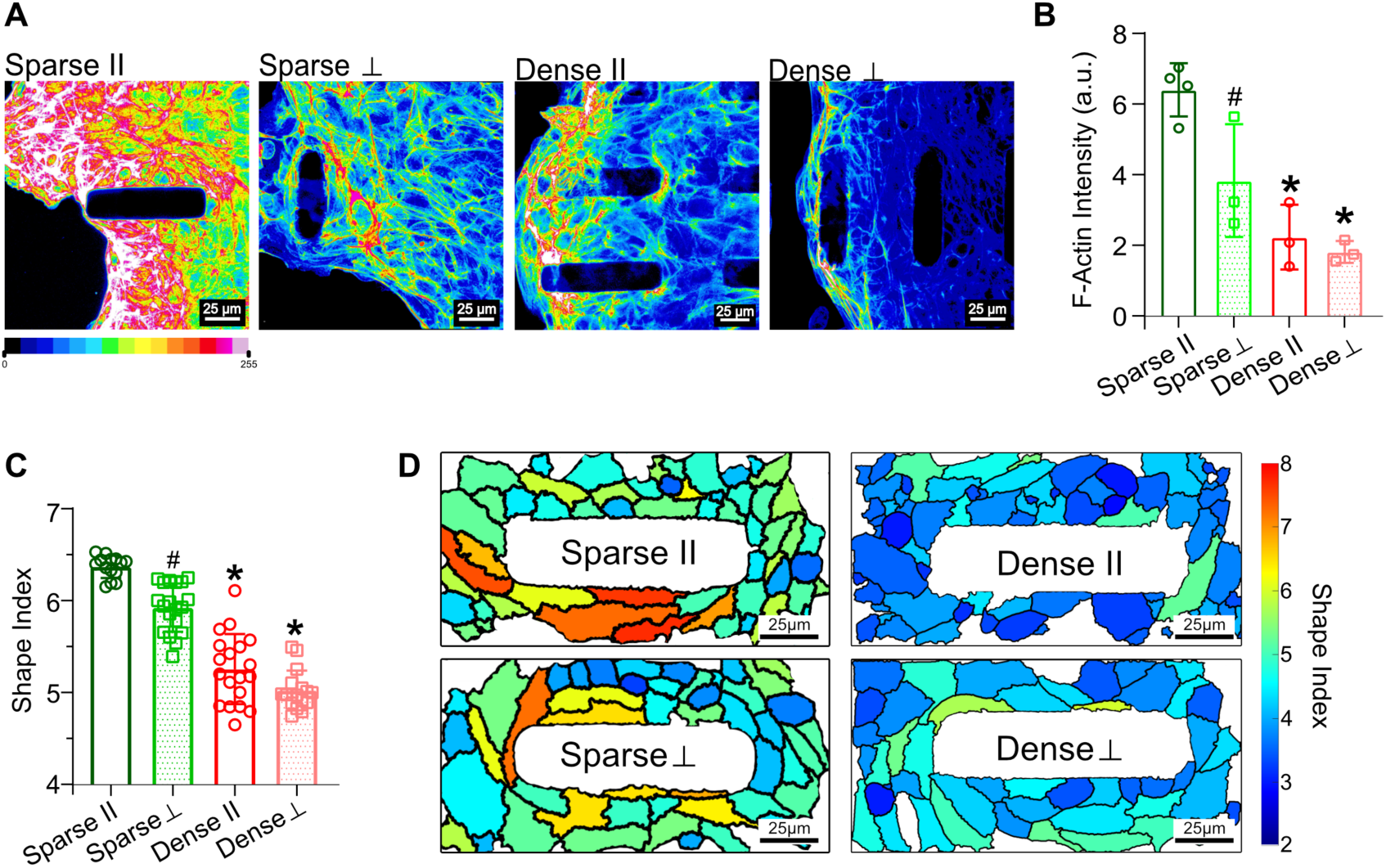
Dense obstructions reduce cell shape fluidity. **(A)** Representative images showing F-actin intensity (phalloidin) in cell monolayer around dense or sparse obstructions with parallel or perpendicular orientations. **(B)** Average F-actin intensity normalized by area occupied by cells across. Error bars = SD, N ≥ 3 samples. **(C)** Average cell shape index and **(D)** representative cell shapes color-coded with cell-level shape index values. Error bars = SD, N ≥ 10 regions of interest around stumps. Significant difference annotated as *p<0.05 for comparison against the sparse condition, and #p<0.05 for comparison against the parallel condition.

In previous work, various collective cell migration phenotypes have been described in terms of jamming and unjamming within epithelial colonies derived from healthy or diseased states, e.g., in asthma and breast cancer (19, 21). In these cases, the ‘unjammed’ cells move in a fluid-like state, characterized by high shape index, defined as 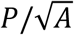 (*P* is the perimeter and *A*is the area of cells that form the colony). In our system, the same healthy epithelial colony alters its migration phenotype, mechanical properties, and cytoskeletal composition due to the presence of matrix obstructions – fast and fluid-like migration on substrates with sparse obstructions changes to more frustrated migration through dense obstructions (Fig. 1). Notably, these multiscale changes in migration occur without any prescribed biochemical, genetic, or pathological alterations in cells. The loss of migration fluidity in dense obstructions was accompanied by physical softening and loss of cytoskeletal actin (Fig. 2, Fig. 3A,B). To understand whether cell shape fluidity follows these trends, we measured the shape index of the cells in epithelial monolayers migrating around the stumps. As plotted in Fig. 3C and visualized in Fig. 3D, the shape index of cells around sparse stumps was higher compared to dense stumps, which indicates that cell shape fluidity reduces due to cell interaction with densely packed obstructions. Here, the cell shape index was the highest in the case of sparse-parallel obstructions, which is consistent with the trends we observed above for cell stiffness and actin intensity.

### Lattice-based modeling to understand obstructed collective cell migration

Physical understanding of collective cell migration has been developed through direct cellular measurements in concert with computational models based on cell-generated forces, cell-cell cooperation, and cell-ECM adhesions (16). Previous models have combined physics-based and phenomenological approaches. For example, cells can be treated as agents that move according to a net energy minimization criterion where the rules of cohesion and scattering are incorporated in terms of inter-cellular potentials (35, 36). In dynamic vertex-based models, cell migration is generated from nodal movements that minimize local energy and fluidic collective motion is captured through dynamic cell shape changes (20). Similarly, lattice-based models generate single and collective cell migration such that any movement minimizes the net system energy (37, 38). In these models, physics-based components form the basis for the model and then simple rules are implemented to capture experimental observation, which leads to new insights regarding how any given biological system differs from the original physical system. Along these lines, to understand how obstructions alter collective cell migration, we first develop a simple lattice-based model for collective cell migration and apply as few rules as possible to capture experiments. To this end, we first note a few experimental observations that had not been shown before this study: (1) dense obstructions induce follower-like cell response in terms of reduced cell stiffness, actin expression, and migration phenotypes, and (2) mesoscale obstructions require cell-cell communication to elicit macroscale response in the form of disordered leading edge and reduced collective migration speed and directionality. We developed a cellular Potts model for collective cell migration in the presence of obstructions (as detailed in Materials & Methods). Briefly, protrusive energy *α* and contractile energy regulated by parameter *ϕ* govern active cell migration energy. In normal situations (without obstructions), leader cells pull on follower cells to generate collective migration.

According to our experiments, it is likely that the protrusive identity of leader cells is disrupted by the obstructions. To capture this, when leader cells come in contact with obstructions, our model assumes that active forces generated by a leader cell are reduced through a rise in contractile energy (via *ϕ*) and reduction in protrusive energy (*α*_*L*_), as described in Fig. 4A. Given the known macroscale response of obstructions on collective migration, we also assume that signals from leader cells are immediately propagated throughout all cells in the monolayer.

**Figure 4:**
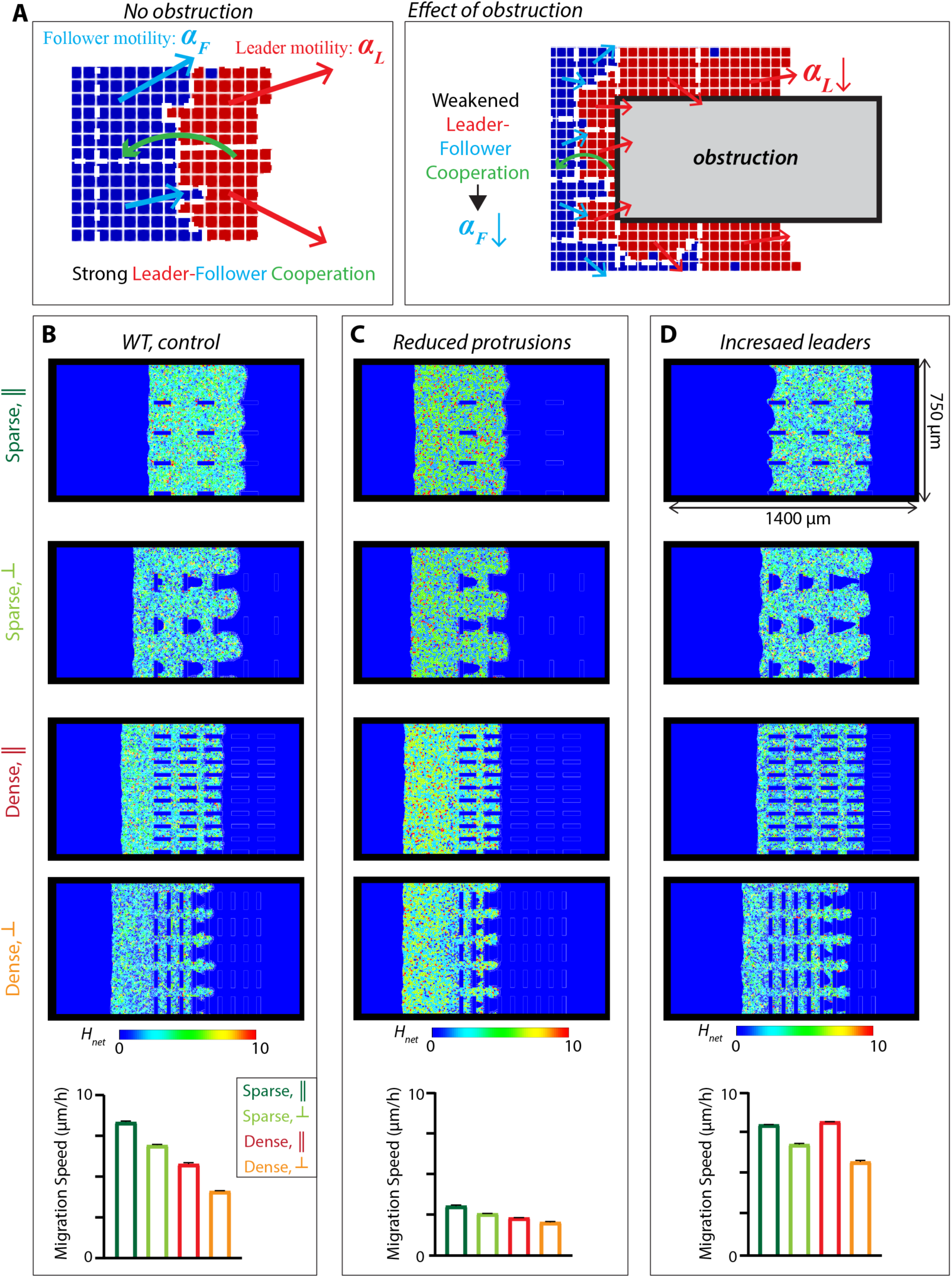
Lattice-based modeling of collective cell migration through matrix obstructions. **(A)** Cellular Potts visualization of a migrating cell colony describing high leader cell motility and leader-follower cooperation (on the left) and weakened leader cell motility and reduced leader-follower cooperation in the presence of obstructions (on the right). **(B)** Simulations are performed and monolayer state is plotted at *t = 54h* (other timepoints are provided in Supplementary Figure 3). Each cell in the colony is color-coded with net energy *H*_*net*_, whose higher values indicate higher energy cost and thus predict reduced resulting migration. Average migration speed after at least 5 simulations is compared in the bottom panel. Simulations and migration speed predictions are repeated for cases with **(C)** reduced protrusions in all cells and **(D)** induction of leader-like properties in all cells.

We simulated a monolayer of about 4000 cells migrating into sparse and dense obstructions in parallel and perpendicular orientations. The front-rear polarity in each cell is randomly chosen in the range of angles between −90° and 90°, with reference 0° angle in the horizontal rightward direction. Thus, the entire monolayer moves forward into the region with obstructions. We performed simulations of cell migration through obstructions over 4000 Monte Carlo Steps (MCS) equivalent to 3 days. All monolayers started at the same point, outside the region with obstructions (Supplementary Fig. 3 showing simulated cell monolayers at different timepoints), and the monolayer at the final timepoint is visualized in Fig. 6B for sparse and dense obstructions with parallel and perpendicular orientations. Here, net monolayer translocation and average migration speed were reduced with obstructions, which is broadly consistent with experiments. As visualized through color-coded cells, the net energy cost (*H*) of cell migration increases with a higher number of obstructions. This occurs due to our model of reduced change in active energy from protrusions and contractility as more leader cells interact with dense obstructions. Also similar to experiments, the leading edge integrity was more compromised in dense obstructions, because of higher heterogeneity in net energy at the leading edge. These simulations indicate that the simple modeling assumption of suppressed leader cell function upon contact with obstructions is enough to capture our experimental observations.

Since individual cell motility is crucial for collective obstruction sensitivity, we next reduced protrusive potential of all cells. In our model, we reduced active energy for all cells to diminish the effect of obstructions on active leader cells, which was implemented by increasing *ϕ* from 0.1 (WT) to 3. Simulations for this case (Fig. 4C) showed reduced differences due to obstruction density and orientation, with overall slower collective migration compared to WT. The spatial plots of net energy showed that slower migration in dense obstructions is associated with the rise in energy cost. Therefore, a protrusion-mediated microscale change leads to reduced ability of cells to sense the obstructions. Next, we explore a reversed situation – if all cells are prescribed to maintain leader-like high protrusions despite their mesoscale contact with obstructions, would the whole cell population be able to ignore obstructions altogether? In this simulation, all cells would behave as leader cells without cell-cell communication. We assumed that all cells have the same protrusive energy as the leader cells (*α*_*F*_ = *α*_*L*_ = 8). These simulations showed an overall faster migration compared to WT (Fig. 4D) and minimal sensitivity to obstruction density and orientation. Overall, our model indicates that the phenotypic suppression of the leader cell upon contact with obstructions is a critical feature of their observed multiscale effect on collective cell migration.

### Rac inhibition reduces but does not eliminate obstruction-sensitive migration

According to our simulations, microscale suppression of protrusive leaders reduces obstruction-sensitive collective migration (Fig. 4C). We have also shown that cells tumbling around dense obstructions lose their shape fluidity, cytoskeletal structure, and stiffness (Figs. 2,3), generating macroscale reduction in cell migration speed and compromised leading edge integrity (Fig. 1). According to our data, cells sense and respond to matrix obstructions on a cytoskeletal level, particularly F-actin expression and cell stiffness. Furthermore, the loss of directionality in dense obstructions indicates that front-rear polarity within the cells may be disrupted as they move around stumps. Given the known role of the small GTPase Rac in actin polymerization and cell polarity in directed cell migration (39, 40), we argued that inhibition of Rac could reduce the ability of epithelial cells to sense the density of obstructions. To test these predictions, we performed pharmacological Rac inhibition by treating cells with NSC23766 and monitoring their migration for 24hrs. From migration videos, we noticed a clear change in migration phenotype through dense obstructions – leading edges consistently advanced over time (Fig. 5A, each time frame visualizations provided in supplementary Fig. S4) without forming chaotic geometries or backtracking that we had observed in the untreated wildtype (WT) case (Fig. 1). Leading edge advancement through sparse obstructions was still higher than through dense obstructions; however, the difference between the two conditions was much less obvious compared to the WT condition. Here, stump orientation did not seem to elicit a clear difference in cell migration phenotypes.

**Figure 5:**
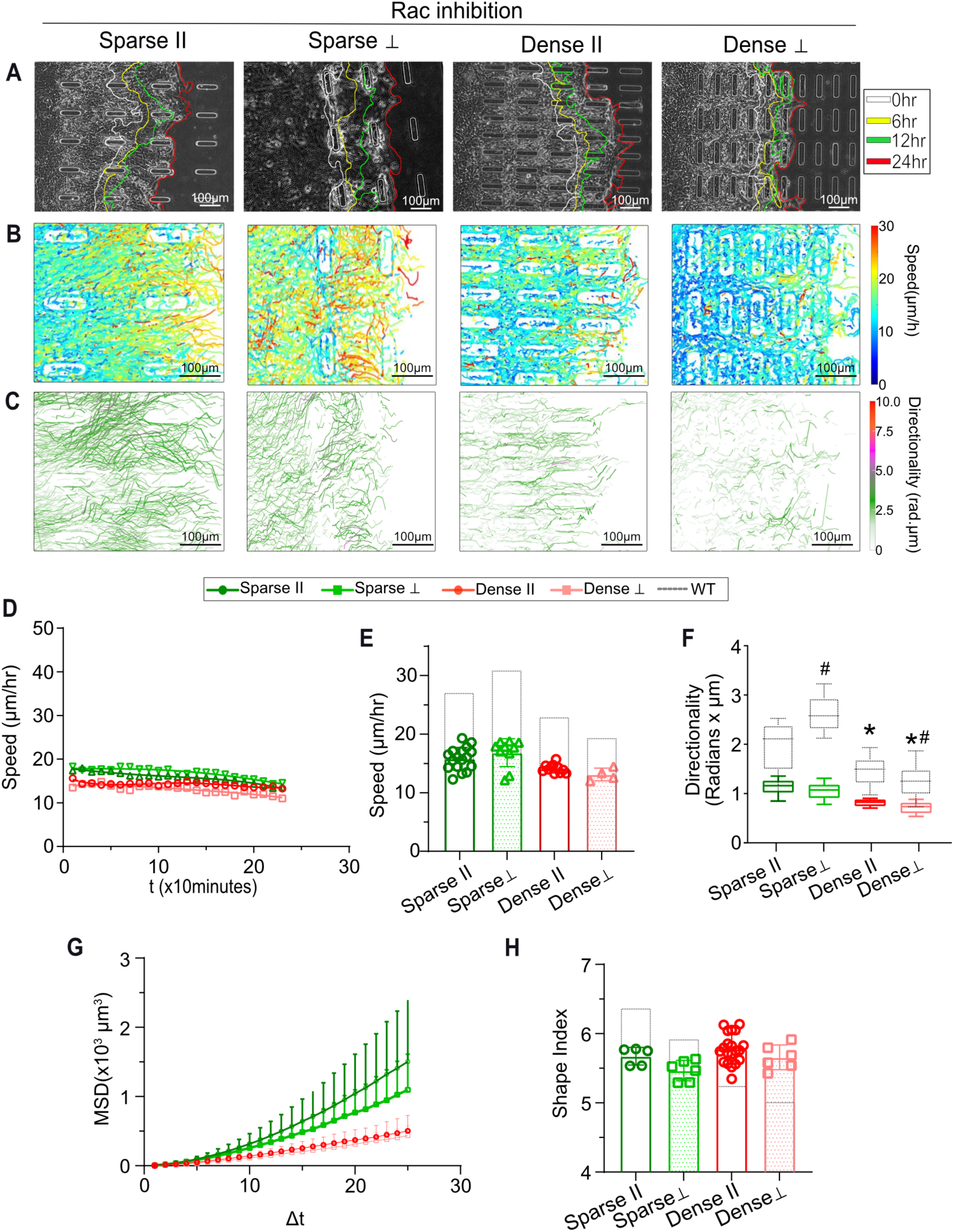
Rac-inhibition reduces obstruction sensitive collective cell migration. **(A)** Representative phase contrast images at final timepoint (24hr after cell seeding) of MCF10A epithelial cells treated with Rac inhibitor in four substrate conditions with varying obstruction spacing and orientation. Leading edges of migrating colonies are annotated for timepoints at 6hr interval between 0-24hr. Migration trajectories of individual cells within Rac-inhibited epithelial monolayers are plotted with color coding for time-averaged **(B)** speed and **(C)** directionality. **(D)** Temporal variation in average migration speed for each substrate condition; points represent average across all cells for a specific timepoint and lines are polynomial fits across timepoints. **(E)** Speed and **(F)** directionality of cell migration averaged over all cells per condition; dotted lines and boxes show data for WT for direct comparisons. Error bars = SD. **(G)** Plot of mean squared displacement (MSD) versus duration of time interval. Error bars = SEM. **(H)** Average cell shape index plotted for all four substrates. Error bars = SEM. Significant difference annotated as *p<0.05 for comparison against the sparse condition, and #p<0.05 for comparison against the parallel condition. N ≥ 6 fields of view in all plots.

To quantify and visualize the spatial distribution of migration speed on a cellular scale, we plotted cell migration trajectories and color-coded them with their average speeds (Fig. 5B). From these speed colormaps, it became clearer that when migrating through sparse obstructions, cells near the leading edge were faster compared to those in the monolayer interior (Fig. 5B; sparse conditions). Thus, long distance communication between leaders and followers may still be occurring on the monolayer scale despite Rac inhibition. By contrast, although cells migrating through dense obstructions were faster compared to WT (with more ‘red’ tracks), those fast cells were somewhat randomly distributed throughout the monolayer, without any clear localization at the leading edge. These observations indicate that in moving through dense obstructions epithelial monolayers lose the sense of leaders and followers. Cell migration speed averaged independently of cell location within the colony stayed consistent over time and was not different between sparse and dense obstructions after Rac inhibition (Fig. 5D). When further averaged across time, although the mean cell migration speed was slightly higher through sparse obstructions, compared to dense, it was not a significant difference (Fig. 5E). Here, we also noted that Rac-inhibited cells became slower compared to WT, with a higher decline in speed on substrates with sparse stumps (Fig. 5E), which is expected given the known role of Rac in cellular protrusions, polarity, and migration (27, 33, 39, 41). Consistent with this trend, the directionality of migration was also significantly reduced compared to WT, rendering it insensitive to obstruction density (Fig. 5C,F). The slight migration advantage through sparse obstructions became more apparent in the relationship between MSD and time interval (Fig. 5G), indicating more fluidic migration. To analyze cell-shape fluidity, we measured shape index in Rac-inhibited cells around the stumps and found no differences across sparse and dense stumps (Fig. 5H). Compared to WT cells, we noted that the shape index decreased around sparse obstructions, but it was rescued in the case of dense obstructions, indicating that the effect of Rac inhibition on cell shape differed for the two cases.

Overall, these results indicate that Rac inhibition reduces the ability of cells to sense and respond to obstructions on the cytoskeletal scale. However, on the macroscale in the whole monolayer, some differences in collective migration phenotypes persisted. In sparse obstructions, there was still a higher migration fluidity (MSD vs time interval in Fig. 5G), and a gradient of increasing migration speed from followers to leaders (Fig. 5B).

### Loss of cell-cell adhesions abrogates macroscale obstruction sensitivity

According to our interpretation of cell stiffness and migration analyses, obstructions make leader cells behave like follower cells. Moreover, cytoskeletal disruption by Rac inhibition did not completely diminish the ability of cells to distinguish sparse and dense obstructions. In our simulations (Fig. 4D), the population-wide activation of leader phenotype (without intercellular cooperation) completely diminishes the effects of matrix obstructions on collective cell migration. To further test obstruction sensing, we sought to disrupt cell-cell communication and alter all epithelial cells towards leader phenotype, which would obviate the possibility of leader-follower communication. We targeted α-catenin, which binds to actin and regulates cell-cell adhesions (42). Since α-catenin is a transcription inhibitor, its depletion enhances YAP activity, induces mesenchymal phenotype, and effectively makes all cells behave like leaders (43, 44).

We used MCF10A cells with depleted α-catenin (10A–αcat) via shRNAi, developed previously (45), and repeated the migration experiments. Across substrates with sparse and dense obstructions, collective cell migration was chaotic, to such a degree that it was difficult to trace leading edges. As visualized in Fig. 6A, although the leading edges of the migrating cell monolayer advanced over time, they were much rougher compared to WT and there was no real difference across sparse or dense obstructions. According to spatial colormaps of migration speed of individual cells, there were no visible leader-follower gradients in speeds, and fast cells (red tracks) were scattered across the monolayer on all substrates (Fig. 6B). This phenotype was particularly unique in the case of dense obstructions, where WT and Rac-inhibited cells were much slower. Indeed, upon averaging cell speeds across multiple samples, cell migration speed was steady over time (Fig. 6D) and insensitive to obstruction density.

**Figure 6:**
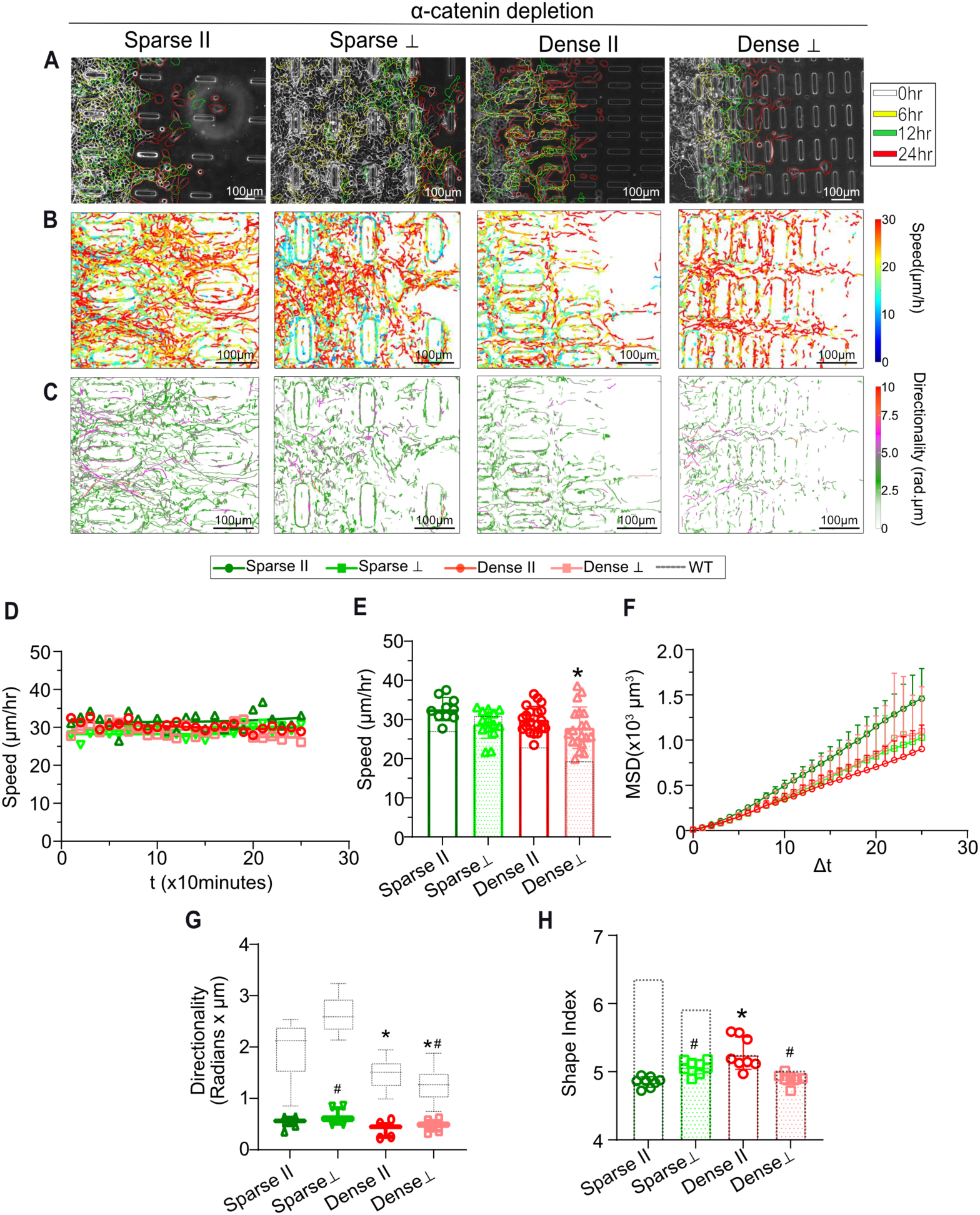
Depletion of α-catenin eliminates obstruction sensitivity in epithelial migration. **(A)** Representative phase contrast images at final timepoint (24hr after cell seeding) of MCF10A with α-catenin-KD (10A–αcat) in four substrate conditions with varying obstruction spacing and orientation. Leading edges of migrating colonies are annotated for timepoints at 6hr interval between 0-24hr. Migration trajectories of individual cells within 10A–αcat cell monolayers are plotted with color coding for time-averaged **(B)** speed and **(C)** directionality. **(D)** Temporal variation in average migration speed for each substrate condition; points represent average across all cells for a specific timepoint, lines are polynomial fits across timepoints. **(E)** Migration speed of 10A–αcat cells averaged over all cells per condition; dotted lines and boxes show data for WT for direct comparisons. Error bars = SD. **(F)** Plot of mean squared displacement (MSD) versus duration of time interval. Error bars = SEM. **(G)** Migration directionality of 10A–αcat cells averaged over all cells per condition. Error bars = SD. **(H)** Average cell shape index plotted for all four substrates. Error bars = SD. Significant difference annotated as *p<0.05 for comparison against the sparse condition, and #p<0.05 for comparison against the parallel condition. N ≥ 6 fields of view in all plots.

Relative to WT cells, although the migration speed of 10A–αcat cells increased on all substrates, this rise was the most substantial in dense obstructions (Fig. 6E), which eventually led to the obstruction-insensitive collective cell migration noted above. The fluidity of 10A–αcat cell migration, assessed by plotting MSD relative to time interval (Fig. 6F), was also insensitive to obstruction density, which was not the case after Rac inhibition (Fig. 6H). While the 10A–αcat cells were, in general, faster compared to WT, their directionality was reduced (Fig. 6G), an indication of random scattering that may explain the chaotic nature of the leading edge (Fig. 6A, each time frame visualizations provided in supplementary Fig. S6). This is not surprising given the loss of cell-cell adhesions after α-catenin depletion, which is expected to abrogate both physical and biochemical signaling communication across the cell colony. Consistent with these migration results, cell shape fluidity measured in terms of shape index was also rendered insensitive to obstruction density (Fig. 6H). Overall, these results indicate that the loss of macroscale cell-cell communication via α-catenin depletion is more potent at abrogating the effects of obstructions on collective cell migration compared to the cell-level disruption via Rac inhibition. These results further indicate that matrix obstructions cause macroscale epithelial response by inducing follower-like behavior across the monolayer. When ‘leader phenotype’ is pre-introduced and cell-cell communication is disabled, obstructions do not have much of an effect on collective cell migration.

## Discussion

Epithelial cell migration is an important process in many biological phenomena, from cellular development to wound repair; and, its disorder can lead to diseased states, such as cancer metastasis and asthma. Due to such biological transformations, collective cell migration has been shown to adopt different modes, described in terms of jamming-unjamming and solid-fluid transitions (19, 20, 26). When cells migrate into external environments, they actively respond to various extracellular physical properties including confinement, stiffness, and surface geometry (7, 9, 27, 46). Cells are also exposed to various topologically diverse environments such as naturally occurring physiological and mechanical heterogeneities in organs, tumors, epithelial folding, or defective basement membranes (28, 47, 48). Adding an ECM perspective to this understanding of collective cell migration, our findings show that topological obstructions in the matrix can trigger physical responses spanning length scales from μm to mm in healthy epithelial cells. With microfabricated stumps on PDMS substrates for epithelial cell culture, we have shown that as the density of the obstructions increased, cells became slower, more disordered, and lost their directional persistence, all of which indicate a less fluidic phenotype. As a result, the leading edge of the epithelial monolayer became more disorganized when moving around dense obstructions. When the spacing between the obstructions (150μm) was larger than their characteristic length (100μm), the overall migration was not significantly impacted. However, migration was more arrested and cell shape fluidity was reduced in dense obstructions, with smaller spacing (50μm). We speculate that dense obstructions cause too many breaks in the epithelial monolayer to retain the original fluidic migration phenotype.

Previous work has connected cell shape fluidity and unjammed collective cell migration to increased tractions generated by diseased cell populations (11, 21). Separately, cell and tissue stiffness have emerged has a ‘mechanotype’ for disease progression – tumor-laden tissues become stiffer (49–51) and cancer cells become more mechanically adaptable depending on their need for invasion (30, 52). Within a given epithelial population, leader cells that generally migrate faster also express higher pMLC and YAP – proteins typically associated with their mechanotransductive activation (9, 27). Through AFM measurements across the distance from the leading edge, we showed that leader cells are significantly stiffer than follower cells within the monolayer. Thus, in addition to traction forces and mechanotransductive protein expression, cell stiffness gradients can also predict leader cell mechanoactivation within a given epithelial population. When moving through dense obstructions, average leader cell stiffness was reduced to the levels observed otherwise for follower cells. Further supported by reduced F-actin expression in dense obstructions, we speculate that obstructed migration may convert leader cells into follower cells in terms of their mechanical properties (Fig. 7A,B). These results show that mesoscale obstructions can cause both microscale and macroscale disruption in cell population – from the shape and stiffness of individual cells to collective migration and leading edge integrity.

**Figure 7.**
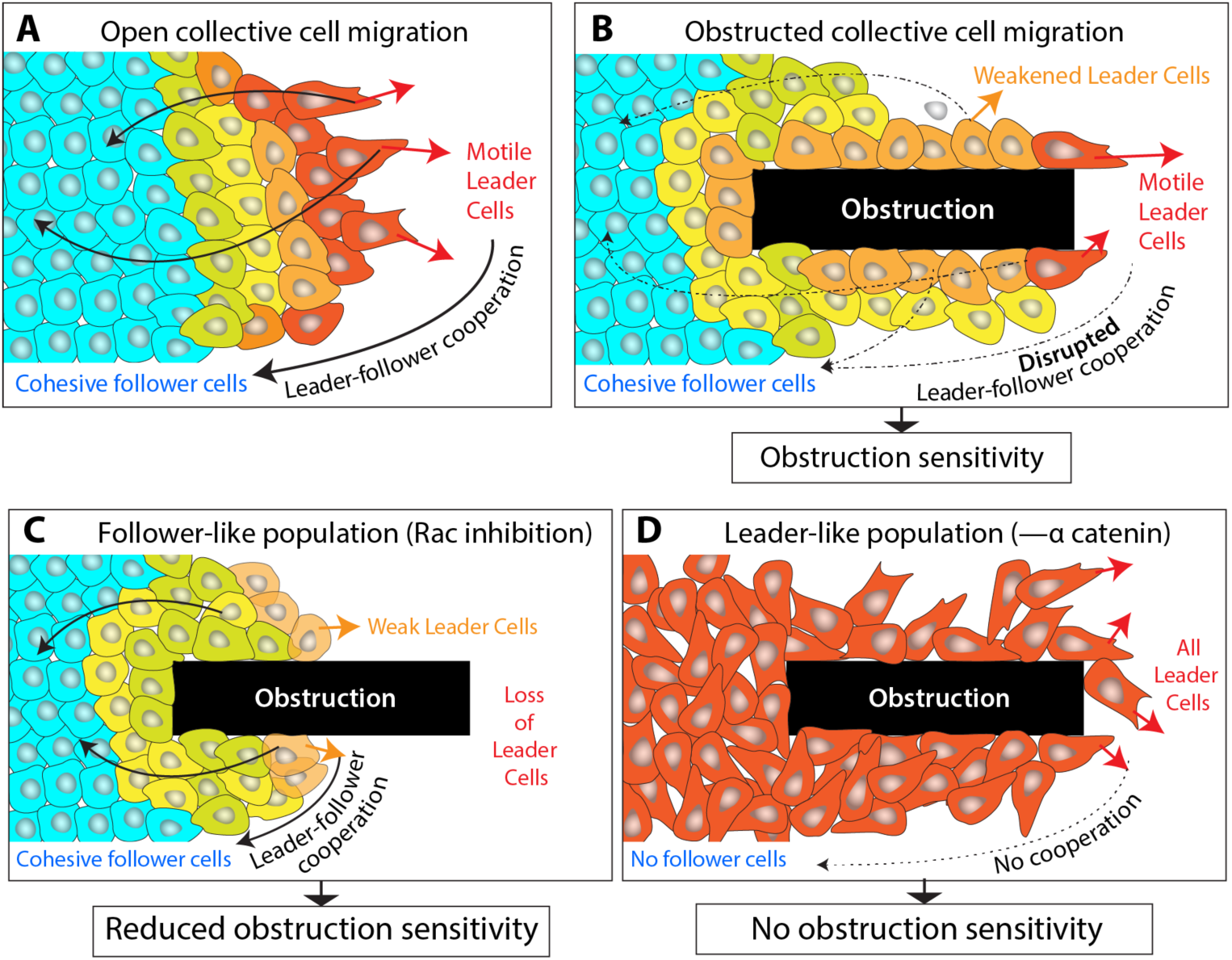
Schematic describing conceptual model for obstruction sensitivity in collective cell migration. **(A)** In unobstructed ‘open’ matrices, motile leader cells pull forward and cooperate with trailing follower cells to enable collective cell migration. **(B)** In the presence of dense obstructions, leader cells are weakened and leader-follower cooperation is disrupted, which slows down collective migration and generates “obstruction sensitivity” due to difference from (A) the open condition. **(C)** When all cells are made to be less motile (cohesive like follower cells) via Rac inhibition, it renders the obstruction-dependent suppression of leader cells pointless, because all cells are suppressed in their motility. As a result, obstruction sensitivity is reduced. **(D)** When all cells are made to be highly motile (aggressive like leader cells), there are no follower cells, causing a loss of obstruction sensitivity.

Based on our AFM measurements, cell shape analyses, and computational modeling, we speculate that the phenotypic leader-follower switching during interactions with dense obstructions may give rise to the ‘obstruction-sensitivity’ shown by healthy epithelial cells. Building on this concept, the obstruction-sensitivity could be targeted by either converting all cells into followers or leaders, thus precluding their phenotypic switching. Indeed, the suppression of the typical protrusive function of leader cells (53, 54), performed here via Rac inhibition, reduces the migratory differences between sparse and dense obstructions (Figs. 5,7C). However, since intercellular communication was still preserved, the obstruction-sensitivity did not vanish. Alternatively, when cell communication throughout the monolayer is suppressed via α-catenin depletion, which effectively made all cells behave like individual leaders, the collective migration became more fluidic, faster, and insensitive to the presence of obstructions (Figs. 6,7D).

Taken together, these findings indicate that cohesive, healthy cell populations are more sensitive to obstructions compared to aggressive, dysfunctional cell populations composed of hyperactive leaders (Fig. 7). This fundamental difference in obstructions sensitivity of cohesive versus aggressive populations may also be applicable in complex collective systems other than cells. Over the past decade, the pathological transformation of cells has been associated with their ability to sense and respond to matrix stiffness (55), detach and leave populations via mesenchymal transition (56, 57), and negotiate confined environments (23, 58). As such, stiffness-sensitivity, cell shape, traction forces, confined migration, and cell stiffness have now been recognized as predictive disease markers for cells (20, 21, 59), which have been used to distinguish them from healthy cell populations and perform drug screening (60, 61). According to our findings, the obstruction-sensitivity of a given cell population may emerge as a disease marker, with lower sensitivity possibly predicting an invasive and scattered disease phenotype. In future work, our integrative methodology of obstruction-sensitive migration analyses, AFM measurements, and computational modeling presented here can be scaled up to test obstruction-sensitivity of a variety of cells and situations in a high throughput manner. Since our method of PDMS-based microfabrication and the resultant substrate design (compatible with standard microscopy) can easily accommodate any number or geometries of obstructions, it can be adapted for different cell types and *ex vivo* tissue samples that may vary in their pertinent length scales of obstruction sensitivity.

## Materials and Methods

### Photolithography for silicon masters with stumps

SU8-2050 photoresist was coated evenly on the silicon master using a spin coater (Brewer Science CEE 200x Spin Coater), and exposed to 365nm conventional UV light. The pattern on the silicon master was in the square with the stumps implanted in the migratory direction. Then the undeveloped photoresist was dissolved using the SU-8 developer solution (MicroChem). The processing time was determined according to the manual provided by Washington University in St Louis cleanroom protocol. Then with a UV-LED mask aligner (KLOE UV-KUB3), photolithography was performed on the silicon master that has been coated. To ensure the dimension of the stumps, the measurement was performed using the KLA Tencor D-100 series profilometer.

### Polydimethylsilane (PDMS) substrate fabrication

Silicon/SU-8 master with the stump design was treated with Trichloro (1H, 1H, 2H, 2H-perfluoroctyl) silane (Millipore Sigma, St. Louis, MO) for 30min inside a vacuum desiccator and dried on a hotplate for 10min at 150°C. SYLGARD-184 silicone-based elastomer polymeric base and curing agent were mixed at a 10:1 ratio, respectively and were poured onto the mold. Then, the PDMS mixture was cured in a vacuum desiccator for 1 hour at RT, after which the temperature was increased to 70°C for 4 hours. Each PDMS substrate was placed in a CL-1000 Ultraviolet Crosslinker with 254 nm bulbs for 10min for sterilization. Surface treatment in preparation for cell culture was accomplished by surface activation in a plasma cleaner (Harrick Plasma, Ithaca, NY) for 5min, after which the PDMS was incubated with a Fibronectin solution at a concentration of 0.05mg/mL overnight at 4°C.

### Cell culture

MCF10A, non-malignant breast epithelial cells, and anti-catenin shRNA (α-cat KD) (both courtesy of Greg Longmore, Washington University School of Medicine) were cultured in DMEM/F12 (Millipore Sigma) composed with 5% Horse Serum (Invitrogen), 20 ng/mL epidermal growth factor (EGF, Miltenyi Biotech Inc.), 0.5 mg/mL Hydrocortisone (Sigma), 100 ng/mL Cholera Toxin (Millipore Sigma), 10 *µ*g/mL Insulin (Millipore Sigma), and 0.2% Normocin (Invitrogen) at 37°C with 5% CO2. Media was changed every 3 days in culture plate. To seed the cell on the microfabricated PDMS, approximately 3-5uL (~20,000 cells) droplet was placed on the dedicated flat area (See supplementary Figure 1A).

### Rac inhibition and α-catenin knockdown

shRNA knockdown of α-catenin was developed by inserting a lentiviral pRLRu vector (62). The inhibition of Rac was performed by directly adding 20μM NSC23766 trihydrochloride to the monolayer of MCF10A. The effect of NSC23766 trihydrochloride was observed to take effect within 1hr after treatment.

### Immunofluorescent staining, imaging, and analysis

Following 3 days in culture, samples were fixed in 4% paraformaldehyde solution for 10min. Samples were washed and treated with 30min of 100nM Phalloidin (Invitrogen) and 10min of 1 ug/mL Hoechst 33342 (Thermo Fisher) in DPBS. Images were taken using the confocal microscope (Zeiss LSM 880; Carl Zeiss Micro Imaging, Germany) with constant acquisition parameters, using a 40x objective with a 2x digital zoom. To prevent bias across conditions, laser intensity and exposure time were kept constant, and at least 2 positions were taken per sample, with at least 3 samples per condition. With acquired Images, the basal area was used to calculate the shape index of each cell through Seedwater Segmenter (63). The shape index was calculated according to the equation:

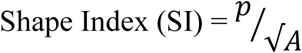

where *p* = perimeter and *A* = area.

### Live Imagining and Real-Time Analysis

Cells (MCF10A or α-Catenin-KD, as applicable) were incubated with 1μg/mL Hoechst 33342 for 10 min, for nuclear labeling, prior to running timelapse imaging. For all conditions, images were taken with a Zeiss AxioObserver microscope (10x objective) equipped with a cell incubator chamber (Carl Zeiss Microscopy) in 10 min intervals for 3 days. The microscope stage incubator was maintained at 37°C with 5% CO_2_ over this time. Images were analyzed with FIJI and the Trackmate plugin (NIH). The raw data was processed through a custom Python (Directionality and MSD), MATLAB (Speed Average and Speed Heatmap), and R (Directionality Heatmap) codes.

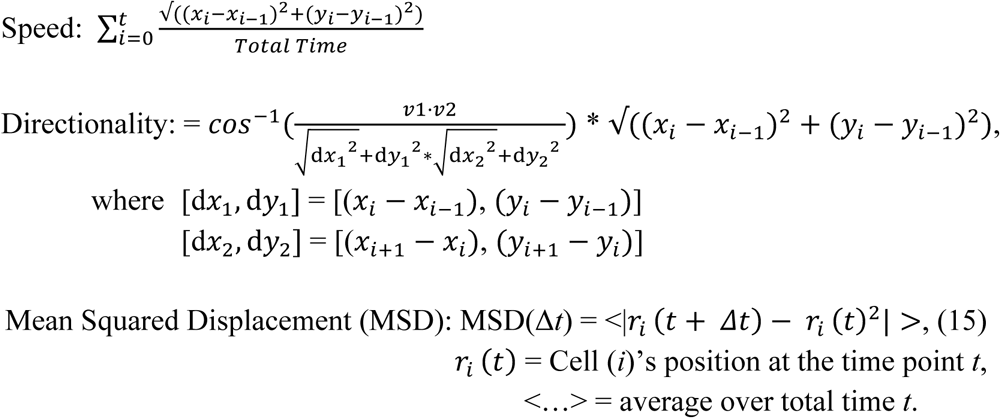

### Atomic Force Microscopy (AFM)

AFM was performed according to previous work (64, 65). All measurements were acquired with a Bruker Nanoscope Resolve atomic force microscope (Bruker, Billerica, USA). Custom AFM probes were used, comprised of a silicon nitride cantilever with a 4.5 μm diameter polystyrene bead attached at the tip (Novascan Technologies, Inc., Boone, IA), with reported stiffness of 0.01 N/m. Tips were initially calibrated in testing buffer to determine the spring constant of the cantilever. After determining the cantilever’s spring constant, the probes were allowed to equilibrate in for at least 10 min to prevent drift in the measured deflection due to the testing buffer. Substrates were loaded onto the AFM stage so that the stumps on the PDMS substrate ran parallel to the AFM cantilever to reduce testing complications from the vertical stumps. Testing began where the leading edges of the monolayers were at the end of a set of stumps. Full force volume measurement runs were taken in 100μm increments until reaching the region of the monolayer before the beginning of the stumps. Each section was taken such that measurements were taken as close to the stumps without the cantilever touching the stumps and thereby skewing the stiffness measurements. A modified Hertz model, implemented in Nanoscope Analysis Software (Bruker, Billerica, USA) was used to analyze the force curves and export stiffness data.

### Statistical Analysis

All box and whisker plot has a representation of mean with 90-10 percentile whiskeys. Statistical significance between groups was determined using one-way ANOVA in GraphPad Prism9 and the p-values <0.05 were considered to be statistically significant, unless specified otherwise.

### Lattice-based modeling of collective cell migration through obstructions

According to our experimental results, obstructions in the path of collectively migrating epithelial cells slow migration speed of individual cells by introducing mesoscale disorder and cell-level softening. Here, we develop a theoretical model and a computational framework to better elucidate these multiscale effects of obstructions on active protrusions and polarity in individual cells and overall intercellular cooperation from leaders to followers within the monolayer. Our goal is to keep the model as simple as possible to capture obstruction-sensitive collective cell migration and predict outcomes as known cellular behaviors are changed. In addition to basic energy-based rules of cellular elasticity and adhesions, we apply two phenomenological effects of obstructions based on experimental observations – upon interaction with obstructions (1) leader cells adopt a follower identity and (2) all cells reduce their active protrusions because active processes cannot go through vertical obstructions.

### Cell migration as an energy minimization problem

The forward motion of a cell within a colony can be understood as dynamic minimization of energy fluctuations due to the temporal evolution of active processes from protrusions and contraction, adhesions with neighboring cells, and passive elastic properties of the cell. Cells would not move if the net change in energy is positive. We define the net change in energy (Fig. 7A) as the sum of energy change required for active protrusions (Δ*H*_*active*_), passive cytoskeletal remodeling (Δ*H*_*passive*_), and formation of cell-cell and cell-ECM adhesions (Δ*H*_*adhesion*_), written as:

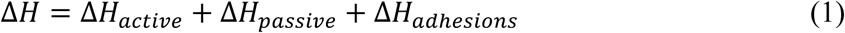

### Adhesion energy

Cellular Potts model (38) represents a space as a discrete collection of lattice points – pixels on a 2D grid or voxels in a 3D grid. We model a collection of biological cells by attaching to each lattice point (*i*; *j*) of a square lattice a label *σ*_*ij*_, which identifies the corresponding cell, and a label *τ*_*ij*_(*σ*_*ij*_), which identifies cell type. Adjacent lattice sites are defined to lie within the same cell if they have the same value of *σ*_*ij*_. System evolves by the random movement of individual pixels that move according to transition probabilities based on Monte Carlo simulations based on the energy criterion described above (66). At each Monte Carlo step, two neighboring pixels are chosen randomly, with one as source pixel and the other as target pixel. If both pixels belong to the same cell (*i. e*., *σ* (*Sourse*) = *σ*(*target*)), then no changes are made to the lattice. Otherwise, the source pixel attempts to occupy the target pixel based on Monte Carlo acceptance probability, which is calculated from the difference in total system energy. The total system energy associated with the configuration, before and after the move, is defined as per Eq. 1. We provide specific definitions for each term in Eq. 1, as following:

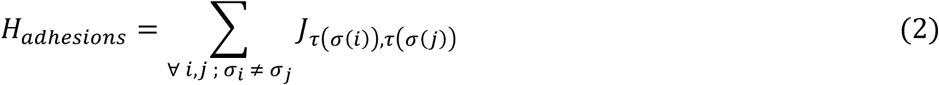

Here, *J*_*τ*(*σ*(*i*))_,_*τ*(*σ*(*j*))_ represents contact energy for the two cell types in contact and (*σ*(*i*)), *τ*(*σ*(*j*)) represents contribution from the total energy due to cell-cell adhesions.

### Passive cytoskeletal energy

Energy due to passive bulk elasticity of cells, *H*_*passive*_, is defined as quadratic change in cell volume, as done previously (38):

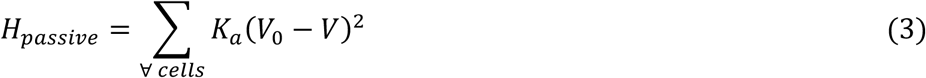

Here, *K*_*a*_ = 5 is a constant for bulk elasticity and *V*_0_ = 200 *µm*^3^ is target volume of the cells in isolation. After calculating energy of system before (*H*_*i*_) and after (*H*_*f*_), the copy attempt will always be successful if *H*_*f*_ < *H*_*i*_, *i. e*. Δ*H* < 0. If Δ*H* ≥ 0, the copy attempt is accepted with a probability of 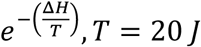. Higher values of Z would tend to accept more unfavorable copy attempt.

### Active protrusive energy and leader-follower cooperation

Cell migrate through coupled processes of forward propulsion via protrusions and active cytoskeletal remodeling. We define net change in energy due to these active processes as:

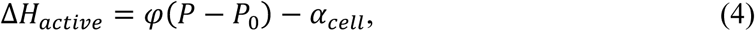

which combines cytoskeletal remodeling (first term) and cell-specific forces (*α*_*cell*_). In the first term, *P* is current cell perimeter and *P*_0_ = 40 *µm* is the target perimeter. As such, higher cell perimeter indicates higher cell spreading, which has to be balanced by lower values of contraction parameter *φ* to enable migration. Higher values of the contraction parameter *φ* result in high energetic cost to extend protrusions. In control conditions, i.e., the open stump-free surfaces, we use *φ*=0.1. Since cells cannot insert protrusions into obstructions, we use *φ*=3 when cells come in contact with obstructions to capture this loss of protrusions.

It has been shown that leader cells generate higher forces and they pull on follower cells via cell-cell adhesions. This leader-follower cooperation due to intercellular cooperation enables collective cell migration. Here, we define leader cells as those at the leading edge of the migrating colony. We define leader cell protrusive energy as *α*_*L*_ = 8, which lowers the net energy to enable migration. For collective migration, leaders must communicate with followers, both mechanically and biochemically. We defined this intercellular communication and cooperation by defining protrusive energy of follower cells as an average of all leader cell protrusions such that follower cell protrusive energy is 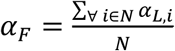, where *N* is the number of leader cells. In the absence of intercellular cooperation, for example in case of α-catenin knockdown, follower cells may also behave as leader cells and thus have the same protrusive energy. Additionally, when leader cells come in contact with obstructions, we assume that they are unable to “lead” migration anymore. Thus, the protrusive energy of leader cells is reduced *α*_*L*_ = 8 to *α*_*L*_ = 4, when they come in contact with obstructions.

### Simulation procedure

We perform cellular Potts simulations within the framework of CompuCell3D and custom Python script to model the cellular response described above. We simulate cell colony over a defined field (700 pixels x 375 pixels, 1px = 2*µm*). Total simulation time is 4800 Monte Carlo Steps (MCSs), equivalent to 72 hours. Cell monolayer is defined as a flat collection of cells of diameter= 10*µm*. Obstructions are defined as areas that cells are not permitted to occupy. Here, we assume that the simulation space is in a 3D space, and show simulations focusing on the basal 2D plane of obstructions and the migrating cell colony.

## Acknowledgements

This work was supported by the NIH/NIGMS MIRA (R35GM128764) grant to AP.

## Author Contributions

AP conceived the project. YLL and CW performed experiments and analyzed data. JM performed computational modeling. HZ contributed software for cell tracking. YLL and AP designed experiments, interpreted findings, made figures, and wrote the manuscript. AP supervised the project and acquired funding.

## References

1. V. Guttal, I. D. Couzin, Social interactions, information use, and the evolution of collective migration. Proc Natl Acad Sci U S A 107, 16172–16177 (2010).

2. C. Blanpain, V. Horsley, E. Fuchs, Epithelial stem cells: turning over new leaves. Cell 128, 445–458 (2007).

3. A. A. Khalil, P. Friedl, Determinants of leader cells in collective cell migration. Integrative Biology 2, 568–568 (2010).

4. J. H. Kim et al., Propulsion and navigation within the advancing monolayer sheet. Nat Mater 12, 856 - 863 (2013).

5. D. T. Tambe et al., Collective cell guidance by cooperative intercellular forces. Nat Mater 10, 469 –475 (2011).

6. T. E. Angelini et al., Glass-like dynamics of collective cell migration. Proceedings of the National Academy of Sciences 108, 4714 –4719 (2011).

7. S. Nasrollahi, A. Pathak, Topographic confinement of epithelial clusters induces epithelial-to-mesenchymal transition in compliant matrices. Sci Rep 6, 18831 (2016).

8. M. H. Kim, Y. Sawada, M. Taya, M. Kino-Oka, Influence of surface topography on the human epithelial cell response to micropatterned substrates with convex and concave architectures. J Biol Eng 8, 13 (2014).

9. M. R. Ng, A. Besser, G. Danuser, J. S. Brugge, Substrate stiffness regulates cadherin-dependent collective migration through myosin-II contractility. J Cell Biol 199, 545 – 563 (2012).

10. N. Hino et al., ERK-Mediated Mechanochemical Waves Direct Collective Cell Polarization. Developmental Cell 53, 646-660.e648 (2020).

11. X. Trepat et al., Physical forces during collective cell migration. Nature Physics 5, 426 – 430 (2009).

12. R. Sunyer et al., Collective cell durotaxis emerges from long-range intercellular force transmission. Science 353, 1157–1161 (2016).

13. V. Te Boekhorst, L. Preziosi, P. Friedl, Plasticity of Cell Migration In Vivo and In Silico. Annu Rev Cell Dev Biol 32, 491–526 (2016).

14. C. De Pascalis, S. Etienne-Manneville, Single and collective cell migration: the mechanics of adhesions. Mol Biol Cell 28, 1833–1846 (2017).

15. J. A. Park et al., Unjamming and cell shape in the asthmatic airway epithelium. Nat Mater 14, 1040–1048 (2015).

16. R. Alert, X. Trepat, Physical Models of Collective Cell Migration. Annual Review of Condensed Matter Physics 11, 77 – 101 (2020).

17. J. A. Park, L. Atia, J. A. Mitchel, J. J. Fredberg, J. P. Butler, Collective migration and cell jamming in asthma, cancer and development. J Cell Sci 129, 3375–3383 (2016).

18. D. Bi, X. Yang, M. C. Marchetti, M. L. Manning, Motility-driven glass and jamming transitions in biological tissues. Phys Rev X 6 (2016).

19. O. Ilina et al., Cell–cell adhesion and 3D matrix confinement determine jamming transitions in breast cancer invasion. Nature Cell Biology 17, 1 – 33 (2020).

20. J. A. Mitchel et al., In primary airway epithelial cells, the unjamming transition is distinct from the epithelial-to-mesenchymal transition. Nat Commun 11, 5053 (2020).

21. J. A. Park et al., Unjamming and cell shape in the asthmatic airway epithelium. Nat Mater 14, 1040 – 1048 (2015).

22. A. Pathak, S. Kumar, Biophysical regulation of tumor cell invasion: moving beyond matrix stiffness. Integrative Biology 3, 267 – 278 (2011).

23. A. Pathak, S. Kumar, Independent regulation of tumor cell migration by matrix stiffness and confinement. Proceedings of the National Academy of Sciences 109, 10334–10339 (2012).

24. R. J. Pelham, Y.-l. Wang, Cell locomotion and focal adhesions are regulated by substrate flexibility. Proceedings of the National Academy of Sciences, USA 94, 13661 – 13665 (1997).

25. S. R. Peyton, A. J. Putnam, Extracellular matrix rigidity governs smooth muscle cell motility in a biphasic fashion. Journal of Cellular Physiology 204, 198 – 209 (2005).

26. X. Trepat, E. Sahai, Mesoscale physical principles of collective cell organization. Nature Physics 13, 58 – 12 (2018).

27. B. Sarker, A. Bagchi, C. Walter, J. Almeida, A. Pathak, Longer collagen fibers trigger multicellular streaming on soft substrates via enhanced forces and cell–cell cooperation. Journal of Cell Science 132, jcs226753 (2019).

28. C. Walter, J. T. Davis, J. Mathur, A. Pathak, Physical defects in basement membrane-mimicking collagen-IV matrices trigger cellular EMT and invasion. Integrative Biology 10.1039/c8ib00034d, 1 – 14 (2018).

29. P. Lu, K. Takai, V. M. Weaver, Z. Werb, Extracellular matrix degradation and remodeling in development and disease. Cold Spring Harb Perspect Biol 3 (2011).

30. C. Rianna, M. Radmacher, S. Kumar, Direct evidence that tumor cells soften when navigating confined spaces. Mol Biol Cell 31, 1726–1734 (2020).

31. B. Sarker, A. Bagchi, C. Walter, J. Almeida, A. Pathak, Longer collagen fibers trigger multicellular streaming on soft substrates via enhanced forces and cell–cell cooperation. Journal of Cell Science 132, jcs226753 (2019).

32. M. L. Gardel et al., Traction stress in focal adhesions correlates biphasically with actin retrograde flow speed. J Cell Biol 183, 999 – 1005 (2008).

33. R. J. Petrie, A. D. Doyle, K. M. Yamada, Random versus directionally persistent cell migration. Nat Rev Mol Cell Biol 10, 538 – 549 (2009).

34. S. Kumar et al., Viscoelastic retraction of single living stress fibers and its impact on cell shape, cytoskeletal organization, and extracellular matrix mechanics. Biophys J 90, 3762 – 3773 (2006).

35. K. Copenhagen et al., Frustration-induced phases in migrating cell clusters. Science Advances 4, eaar8483 – 8410 (2018).

36. B. A. Camley, W. J. Rappel, Physical models of collective cell motility: from cell to tissue. Journal of Physics D: Applied Physics 50, 113002 – 113022 (2017).

37. I. Fortuna et al., CompuCell3D Simulations Reproduce Mesenchymal Cell Migration on Flat Substrates. Biophysj 118, 2801 – 2815 (2020).

38. F. Graner, J. A. Glazier, Simulation of biological cell sorting using a two-dimensional extended Potts model. Phys Rev Lett 69, 2013–2016 (1992).

39. R. Pankov et al., A Rac switch regulates random versus directionally persistent cell migration. J Cell Biol 170, 793 – 802 (2005).

40. A. J. Ridley et al., Cell Migration: Integrating Signals from Front to Back. Science 302, 1704 – 1709 (2003).

41. A. Pathak, S. Kumar, Transforming potential and matrix stiffness co-regulate confinement sensitivity of tumor cell migration. Integrative Biology 5, 1067 – 1075 (2013).

42. B. S. Robinson, K. H. Moberg, Cell-Cell Junctions: &alpha;-Catenin and E-Cadherin Help Fence In Yap1. Curr Biol 21, R890 – R892 (2011).

43. R. L. Daugherty et al., α-Catenin is an inhibitor of transcription. Proceedings of the National Academy of Sciences 111, 5260 – 5265 (2014).

44. A. Vite, C. Zhang, R. Yi, S. Emms, G. L. Radice, α-Catenin-dependent cytoskeletal tension controls Yap activity in the heart. Development 145, dev149823 – 149821 (2018).

45. A. J. Loza et al., Cell density and actomyosin contractility control the organization of migrating collectives within an epithelium. Mol Biol Cell 27, 3459 – 3470 (2016).

46. S. R. Vedula et al., Emerging modes of collective cell migration induced by geometrical constraints. Proceedings of the National Academy of Sciences 109, 12974 – 12979 (2012).

47. C. Pérez-González et al., Mechanical compartmentalization of the intestinal organoid enables crypt folding and collective cell migration. Nature Cell Biology 23, 745–757 (2021).

48. M. Takeda, M. M. Sami, Y.-C. Wang, A homeostatic apical microtubule network shortens cells for epithelial folding via a basal polarity shift. Nature Cell Biology 20, 1 – 15 (2017).

49. C. Walter et al., Increased Tissue Stiffness in Tumors from Mice with Neurofibromatosis-1 Optic Glioma. Biophysical Journal 112, 1535 – 1538 (2017).

50. I. Acerbi et al., Human breast cancer invasion and aggression correlates with ECM stiffening and immune cell infiltration. Integr Biol (Camb) 7, 1120 – 1134 (2015).

51. S. V. H. Bayer et al., DDR2 controls breast tumor stiffness and metastasis by regulating Integrin mediated mechanotransduction in CAFs. Elife 8 (2019).

52. T. H. Kim et al., Cancer cells become less deformable and more invasive with activation of beta-adrenergic signaling. J. Cell Sci. 129, 4563 – 4575 (2016).

53. E. L. Zoeller et al., Genetic heterogeneity within collective invasion packs drives leader and follower cell phenotypes. J. Cell Sci. 132, jcs231514 – 231516 (2019).

54. M. L. Borgne-Rochet et al., P-cadherin-induced decorin secretion is required for collagen fiber alignment and directional collective cell migration. J. Cell Sci. 132, jcs233189 – 233116 (2019).

55. M. J. Paszek et al., Tensional homeostasis and the malignant phenotype. Cancer Cell 8, 241 – 254 (2005).

56. V. J. Guen et al., EMT programs promote basal mammary stem cell and tumor-initiating cell stemness by inducing primary ciliogenesis and Hedgehog signaling. Proceedings of the National Academy of Sciences 114, E10532 – E10539 (2017).

57. D. Hanahan, R. A. Weinberg, Hallmarks of cancer: the next generation. Cell 144, 646 – 674 (2011).

58. C. Beadle et al., The Role of Myosin II in Glioma Invasion of the Brain. Mol Biol Cell 19, 3357 – 3368 (2008).

59. P. S. O. C. Network et al., A physical sciences network characterization of non-tumorigenic and metastatic cells. Sci Rep-uk 3, 1449 (2013).

60. J. G. Lin et al., Linking invasive motility to protein expression in single tumor cells. Lab Chip 18, 371–384 (2018).

61. M. C. Lampi, C.A. Reinhart-King, Targeting extracellular matrix stiffness to attenuate disease: From molecular mechanisms to clinical trials. Sci Transl Med 10 (2018).

62. A. J. Loza et al., Cell density and actomyosin contractility control the organization of migrating collectives within an epithelium. Molecular biology of the cell 27, 3459–3470 (2016).

63. D. N. Mashburn, H. E. Lynch, X. Ma, M. S. Hutson, Enabling user-guided segmentation and tracking of surface-labeled cells in time-lapse image sets of living tissues. Cytometry A 81, 409–418 (2012).

64. B. Sarker, A. Bagchi, C. Walter, J. Almeida, A. A.-O. Pathak, Longer collagen fibers trigger multicellular streaming on soft substrates via enhanced forces and cell-cell cooperation. LID - 10.1242/jcs.226753 [doi] LID - jcs226753.

65. S. Nasrollahi et al., Past matrix stiffness primes epithelial cells and regulates their future collective migration through a mechanical memory.

66. M. H. Swat et al., Multi-scale modeling of tissues using CompuCell3D. Methods Cell Biol 110, 325–366 (2012).

