## Supplementary Information for "Matrix obstructions cause multiscale disruption in collective epithelial migration by suppressing physical function of leader cells"

### Supplementary Figures

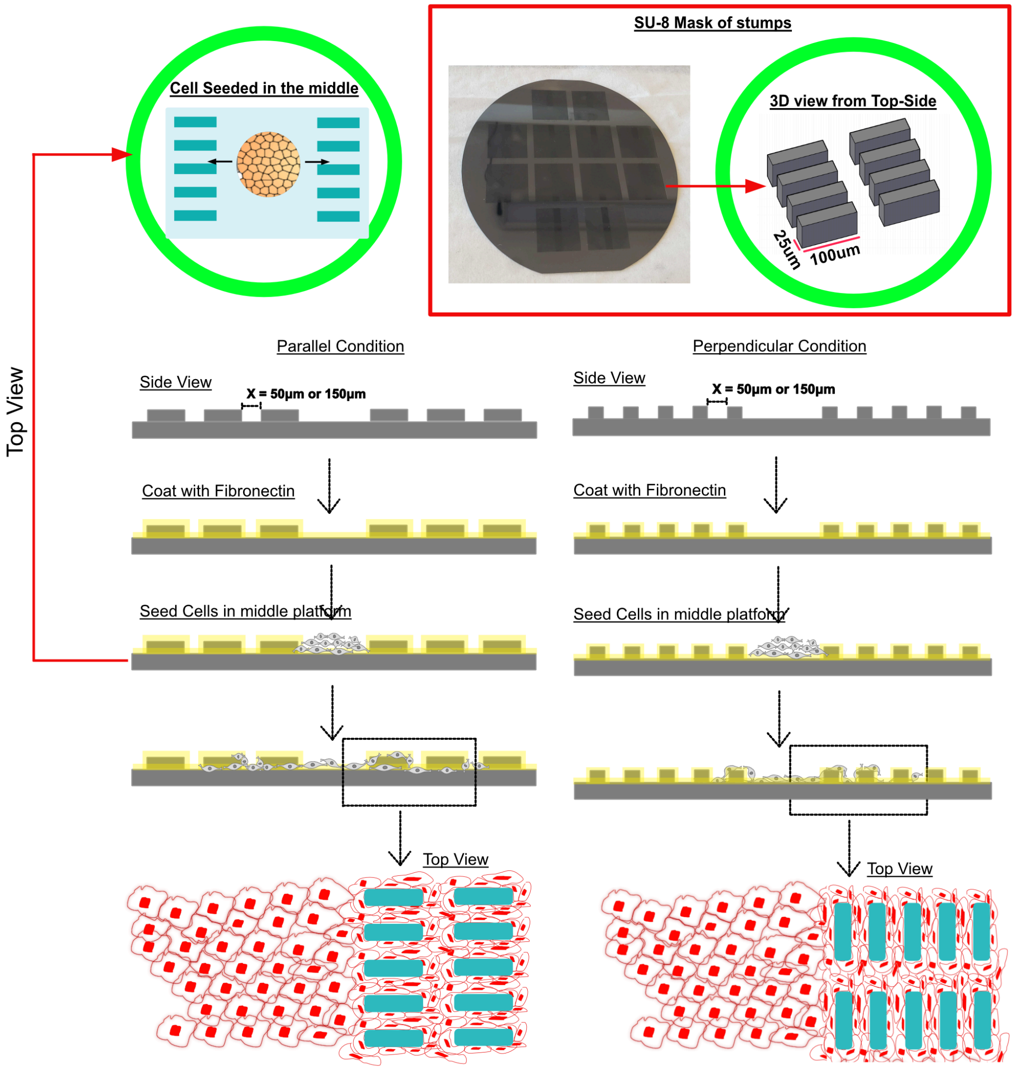


**Supplementary Figure 1. Lithography steps for PDMS substrate preparation.** Using hard lithography, a SU-8 mold is fabricated on silicon discs. These masters are used to perform soft lithography and microfabricate PDMS substrates with defined geometry for stumps. Cells are seeded in the middle and outward collective cell migration is analyzed.

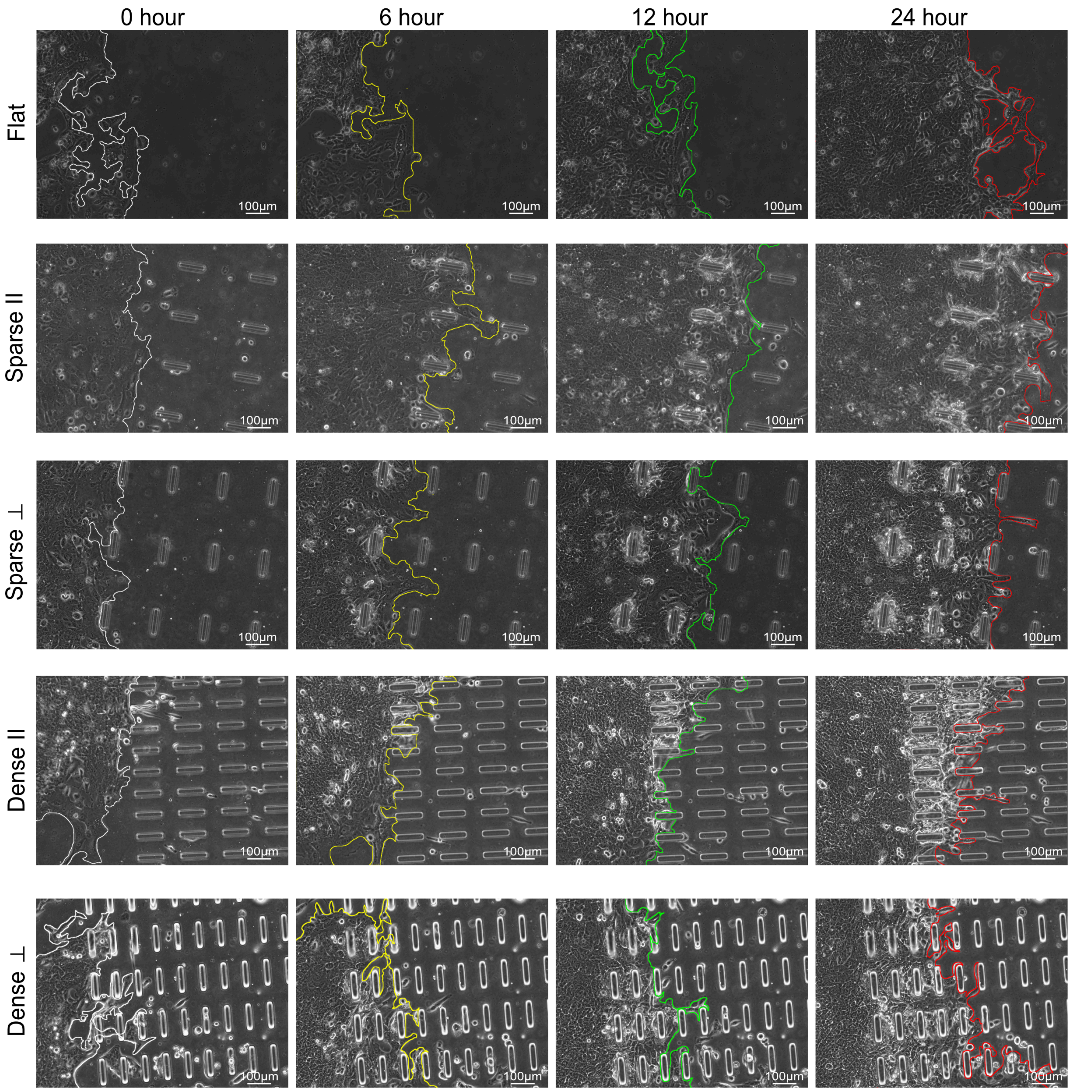


**Supplementary Figure 2. Leading edge shapes for MCF10A epithelial monolayers.** Representative phase contrast images at different timepoints between 0-24hr at 6hr interval, with annotated leading edge shapes, for 5 substrate conditions of varying stump spacing and orientation. Scale bar = 100µm.

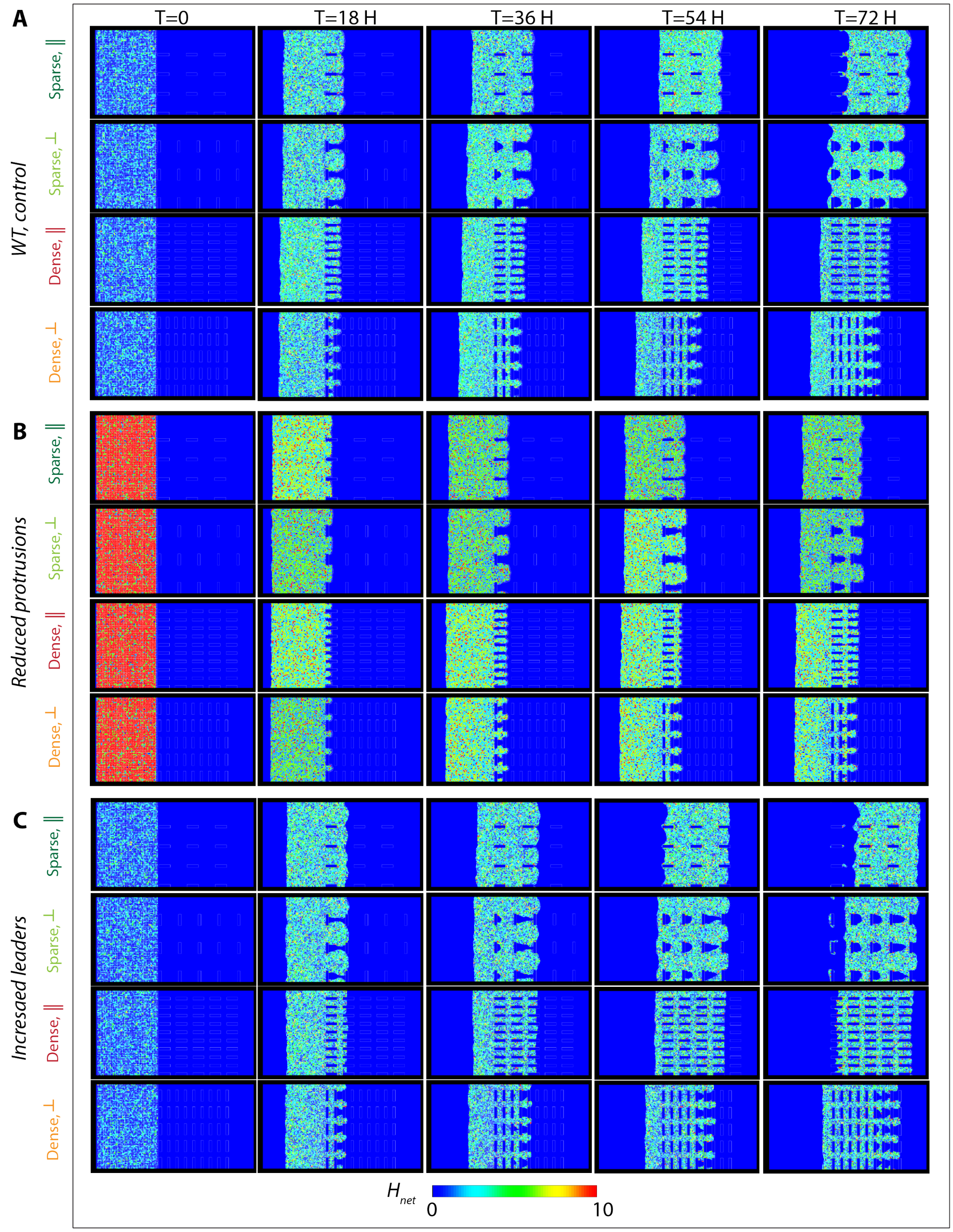


**Supplementary Figure 3. Temporal progression of collectively migrating cells through different obstructions and cell-intrinsic properties.** Snapshots of simulated cell monolayer at different timepoint $(t = 0, 18, 36, 54\ and\ 72\ H)$ for **(A)** the control/WT condition, **(B)** after reduced protrusions, and **(C)** increased leader-like protrusions and lost cell-cell communication. Color coding represents normalized net energy cost.

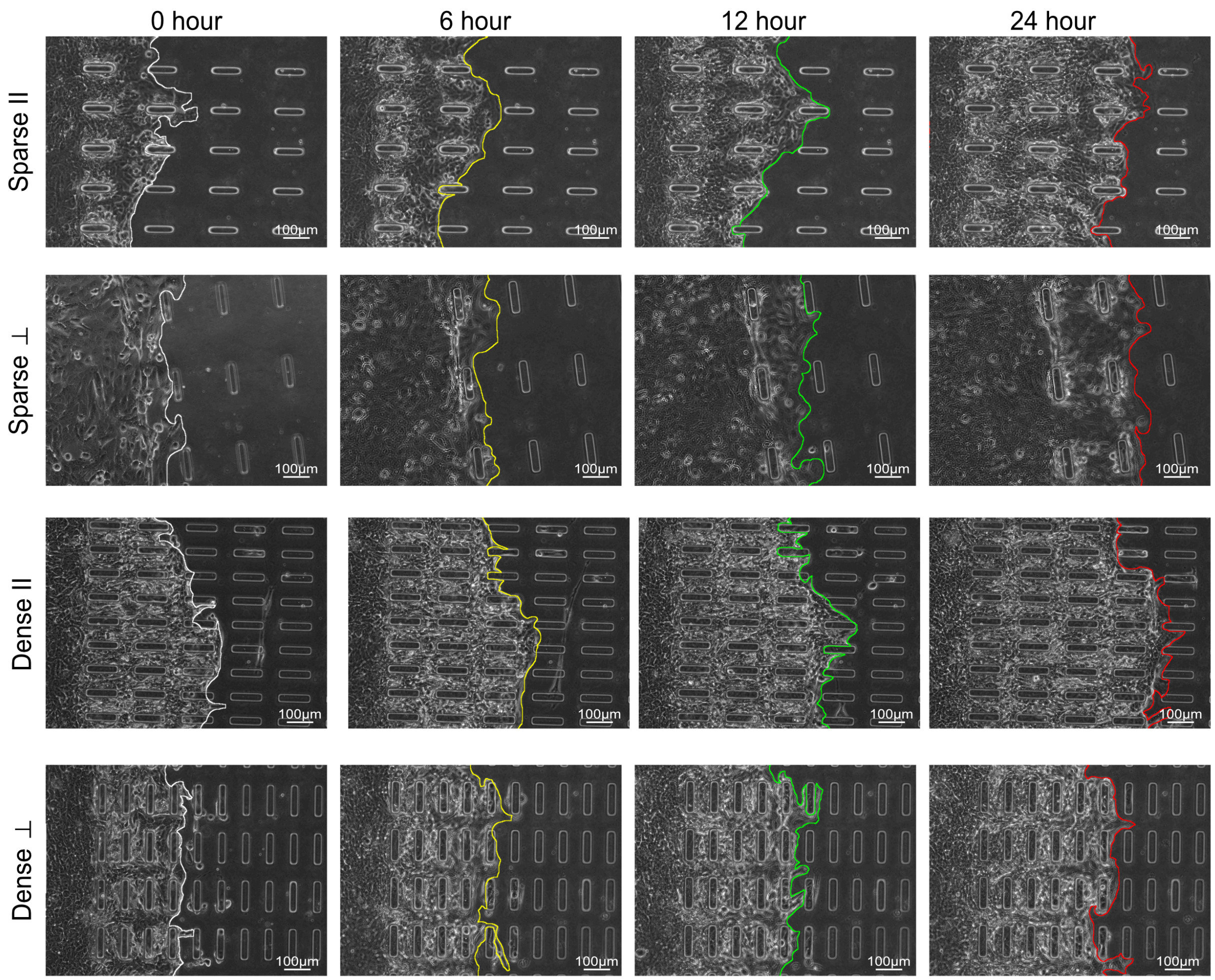


**Supplementary Figure 4. Leading edge shapes for MCF10A epithelial monolayers after Rac inhibition.** Representative phase contrast images of collectively migrating MCF10A cells treated with Rac inhibitor at different timepoints between 0-24hr at 6hr interval, with annotated leading edge shapes, for 4 substrate conditions of varying stump spacing and orientation. Scale bar = 100µm.

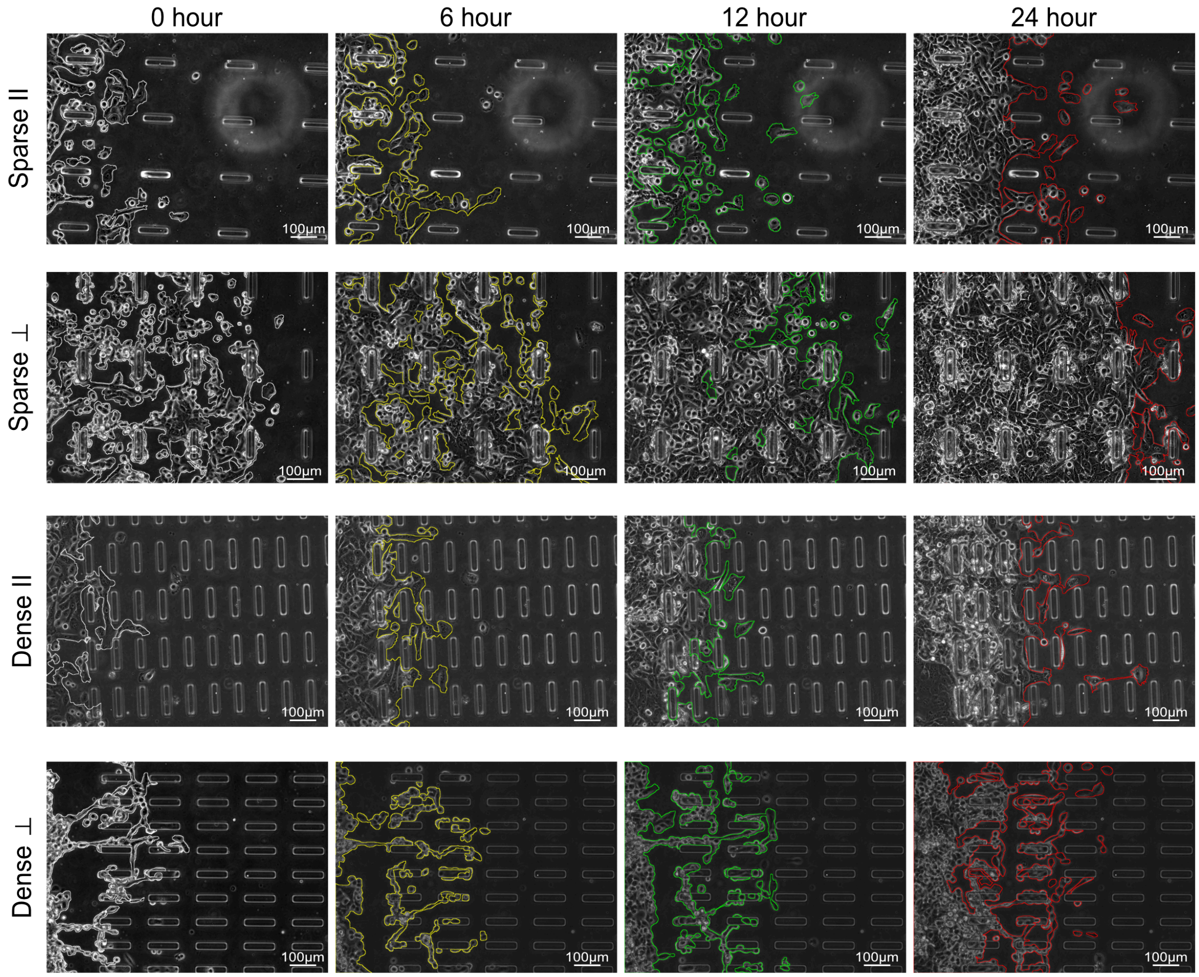


**Supplementary Figure 5. Leading edge shapes for MCF10A-α-Catenin-KD epithelial monolayers.** Representative phase contrast images of collectively migrating MCF10A cells with depleted α-catenin at different timepoints between 0-24hr at 6hr interval, with annotated leading edge shapes, for 4 substrate conditions of varying stump spacing and orientation. Scale bar = 100µm.
